# Structural Metrics from Predicted Protein-Ligand Complexes Enable OATP Inhibitor Classification

**DOI:** 10.64898/2026.11.24.768978

**Authors:** Logan Garland, Ailin Patimar, Alexander Aljets, Daniel Woldring

## Abstract

Organic anion transporting polypeptides (OATPs) mediate the cellular uptake of many clinically important drugs, and their inhibition can alter drug disposition and contribute to drug-drug interactions. Recent advances in protein-ligand co-folding have expanded opportunities for modeling transporters and their interactions with diverse small molecules. However, the extent to which these models can predict functional transporter inhibition remains unclear. Here, we systematically evaluated the predictive utility of structural metrics generated by AlphaFold3, Boltz-2, Chai-1, and Protenix using experimental inhibition measurements for 224 compounds across OATP1B1, OATP1B3, and OATP2B1. Individual structural metrics provided substantial inhibitor discrimination ability, reaching an ROC-AUC of 0.888 for Chai-1 structures of OATP1B3. We then investigated whether integrating multiple structural descriptors improves inhibitor classification relative to optimized individual metrics, which improved performances in many model-OATP combinations. Following combined-metric analysis, Protenix exhibited both the highest absolute average precision (AP) and the largest improvement in AP overall for OATP1B1 (0.560 to 0.763). Mean improvements persisted across five independently generated cross-validation partitions. Performance and the benefits of metric integration differed across transporters and modeling pipelines, while increasing structural ensemble size for each drug from 1 to 100 provided varied improvements. Together, these findings establish the utility of co-folding structural metrics as functional classification features, and this work establishes a framework for evaluating and integrating structure-prediction outputs in broader functional studies of membrane transporters.

## Introduction

Organic anion transporting polypeptides (OATPs) mediate the cellular uptake of structurally diverse endogenous compounds, xenobiotics, and drugs.^1–4^ OATP1B1 and OATP1B3 are predominantly expressed in hepatocytes and contribute to hepatic drug disposition, whereas OATP2B1 has a broader tissue distribution that includes the liver and intestine.^2–5^ These transporters facilitate the uptake of clinically important drugs such as statins into hepatocytes for metabolism or elimination.^6–9^ Inhibition of hepatic OATP uptake can reduce drug clearance, increase systemic exposure, and contribute to clinically significant drug-drug interactions (DDIs). FDA and EMA guidelines recommend evaluating the OATP1B1 and OATP1B3 inhibition potential of investigational drugs during development.^10–13^ Identifying compounds that inhibit OATP-mediated uptake is therefore an important component of early drug development and evaluating potential DDI risks.

Structural characterization of OATPs has historically been challenging due to their flexible and membrane-embedded architecture. Cryo-electron microscopy structures of OATP1B1 and OATP1B3 reported in 2023 confirmed previously suspected transporter conformations and ligand-recognition regions, providing a new experimental foundation for structure-based modeling.^14,15^ However, the new availability of experimentally resolved OATP structures does not mean that OATP function is fully understood. These structures capture selected conformational states rather than the complete transport cycle.^4^ Ligand recognition and transport are linked to significant conformational changes, thus complicating attempts to determine transport behavior from a single modeled protein-ligand complex.^14–16^

Predicting functional inhibition is a very different challenge from predicting protein-ligand binding. Inhibition assays measure changes in the uptake of a probe substrate in the presence of a test compound and provide a metric of transport activity rather than affinity.^17–20^ Reported inhibitory potency can vary substantially across experimental conditions, including probe substrate selection, cell-expression systems, and incubation conditions, resulting in markedly different IC50 values for the same inhibitor.^18–23^ This variability complicates the integration of different inhibition measurements from independent studies. To minimize these inconsistencies, we restricted our experimental benchmark to a single study, in which compounds were evaluated at common test and probe concentrations using consistent uptake assay conditions.^17^ This approach provides a more internally consistent basis for inhibitor classification by limiting variability introduced by different experimental methodologies.

Recent advances in protein-ligand structure prediction have substantially expanded the range of computational tools available for studying these interactions. AlphaFold3, Boltz-2, Chai-1, and Protenix provide complementary implementations of this approach and report model-specific confidence, uncertainty, interface, and ranking metrics alongside predicted structures.^24–27^ These measurements describe different aspects of the predicted complex, but they were not specifically developed to predict function in any capacity.^24,28,29^ Therefore, the relationship between individual structure-prediction metrics and biological function such as transporter inhibition remains poorly understood.

Previous OATP inhibition models have relied on ligand physicochemical properties and molecular representations, with more recent approaches using docking-derived protein interactions for inhibition prediction.^30–36^ A complementary strategy is to test the numerical metrics reported directly by modern co-folding pipelines. Even a single descriptor may capture an experimentally useful inhibition signal, despite not having been designed to associate with transport function. Combining descriptors could provide additional information when individual metrics measure complementary aspects of a predicted complex, although correlated or uninformative features may limit the benefits of metric integration.

Here, we evaluated whether co-folding model structural metrics can distinguish experimentally classified OATP inhibitors from non-inhibitors, and whether integrating multiple structural metrics improves classifier performance. We first benchmarked structural metrics derived from AlphaFold3, Boltz-2, Chai-1, and Protenix against OATP1B1, OATP1B3, and OATP2B1 inhibition measurements from Karlgren et al.^17,24–27^ These were used to establish a binary inhibitor-classification benchmark, with compounds classified according to a 50% transport-inhibition threshold. Single- and combined-metric classifiers were evaluated using compound-grouped nested cross-validation, repeated fold assignments, and randomized-label controls. We further evaluated the predictive contribution of additional predicted complexes across ensemble depths of 1-100 structures and examined descriptor-family ablations to assess predictive robustness. This work establishes the functional predictive utility of single and combined co-folding metrics for OATP inhibition and defines the predictive value and current limitations of co-folding-derived structural information for broader membrane transporter functional studies.

## Methods

### 3.1 Experimental Inhibition Dataset and Classification Endpoint

Experimental inhibition measurements were obtained from Karlgren et al. for 225 compounds tested against the membrane transporters OATP1B1, OATP1B3, and OATP2B1.^17^ The reported measurements describe changes in model-substrate uptake following exposure to each test compound.^18,20^ Because these assays measure functional transport inhibition rather than direct protein-ligand binding affinity, the experimental values were used to establish a binary inhibitor-classification rather than a continuous prediction.

Compounds producing ≥50% inhibition were classified as inhibitors for the corresponding OATP isoform; compounds below this threshold were classified as non-inhibitors. The 50% threshold serves as a clear operational definition of inhibition under the tested assay conditions and is not intended as a validated clinical drug-drug interaction threshold. Each compound was evaluated separately for each transporter: 224 compounds tested against three OATP isoforms yielded 672 compound-isoform observations. Oxaliplatin was excluded from the original 225-compound collection because its platinum-containing chemistry was incompatible with the Boltz-2 prediction workflow. The final benchmark contained 170 inhibitor observations and 502 non-inhibitor observations, including 78 OATP1B1 inhibitors and 46 inhibitors each for OATP1B3 and OATP2B1. Across the 224 compounds, 133 were defined as noninhibitors for all three OATP isoforms. Dataset composition and class balance are summarized in Supplementary Table S1, while experimental inhibition distributions are shown in Supplementary Figure S1. Sensitivity to alternative inhibition thresholds between 25% and 50% was evaluated separately (Supplementary Figure S2).

### 3.2 Protein-Ligand Structure Prediction

Predicted OATP-ligand complexes were generated using four contemporary protein-ligand structure-prediction pipelines: AlphaFold3, Boltz-2, Chai-1, and Protenix.^24–27,37,38^ Model-specific inputs were prepared from the curated ligand structures and corresponding OATP protein sequences using a unified computational workflow. All four prediction pipelines were run using their standard inference configurations, without modifications to their underlying prediction procedures. Separate software environments and input-generation procedures were maintained to accommodate differences among the prediction tools, while compound identities, OATP sequences, and downstream output organization were kept consistent. For each compound-isoform pairing, an ensemble of 100 structural predictions was assembled for each pipeline, generated across 20 seeds of 5 predictions each. Predictions were generated using model-compatible sampling procedures and deterministic seed assignments. The final dataset contained 672 complete compound-isoform ensembles for each of the four prediction pipelines.

Predicted complexes were ordered using each model’s native ranking criterion: ranking score for AlphaFold3 and Protenix, confidence score for Boltz-2, and aggregate score for Chai-1.^24–27,37,38^ The highest-ranked complex among the 100 predictions from each ensemble was used for the primary classification analysis, based on the native ranking scores for each of the four pipelines. For the subsequent ensemble-depth analysis, the highest-ranked prediction from each seed was included first (20 seeds, 5 poses each), followed by the next highest-ranked prediction from each seed. Ranking scores were interpreted within their own pipelines, not compared numerically across pipelines. Seeded pose grouping is not direct evidence of distinct binding conformations.

Prediction metadata were processed to extract model-derived structural descriptors, including confidence, uncertainty, and ranking metrics. The classification analyses reported here used these prediction-derived numerical outputs rather than independently computed docking scores or explicit interaction fingerprints. Classifier performance was assessed using cross-validated predicted inhibitor probabilities, with receiver operating characteristic area under the curve (ROC-AUC) and average precision (AP) as the primary performance metrics. AP summarizes precision across recall levels and has a prevalence-dependent random-ranking baseline; ROC-AUC quantifies global inhibitor to non-inhibitor discrimination, with a random baseline of 0.5. All structure prediction, pose processing, feature extraction, interaction scoring, and classifier analyses were performed using reproducible Python workflows on the MSU High-Performance Computing Center. Separate software environments were maintained for each structure-prediction and scoring pipeline to preserve model-specific dependencies.

### 3.3 Model-Derived Structural Metrics

Numerical descriptors were extracted from the output metadata of each predicted OATP-ligand complex. These metrics included model-reported measures of structural confidence, predicted interface confidence, structural uncertainty, complex ranking, and predicted steric clashes or disorder. These measures describe properties of predicted complexes and are not direct measurements of transport inhibition.^24–27,37,38^

The primary analysis included 39 structural metrics for AlphaFold3, 15 for Boltz-2, 31 for Chai-1, and 74 for Protenix. These counts represent candidate predictors before training-fold-specific filtering. Administrative metadata describing prediction bookkeeping, including random seeds, sample identifiers, recycle counts, and row indices, were excluded to prevent classifiers from learning information unrelated to the predicted molecular interaction. Candidate metrics were held consistent across ensemble depths for the fixed-depth analysis. A complete inventory of the candidate structural descriptors is provided in Supplementary Table S2.

Within every training partition, features were retained if at least 90% of training values were observed and the feature had nonzero training variance. Missing values were replaced with the corresponding training-set median, and features were standardized using training-set means and standard deviations (scikit-learn StandardScaler). These transformations were fitted separately within each inner training fold and refitted on the full outer training partition before prediction, without using held-out feature distributions or labels.

### 3.4 Individual and Combined Structural-Metric Classification

To test whether structural metrics contain predictive signal for OATP inhibitor classification, we compared two approaches within each structure-prediction pipeline and OATP isoform.

The **individual-metric approach** used a single model-derived descriptor as the input to a classifier. Candidate metrics were evaluated independently, with the metric and classifier hyperparameters selected using training data only. This provided an optimized individual-metric comparator rather than an arbitrary choice of a commonly reported confidence or ranking score.

The **combined-metric approach** used all eligible descriptors from the corresponding prediction pipeline as candidate inputs to a single classifier, with training-fold filtering applied as described above. Only the regularization parameter was selected in the inner folds. L2 regularization permitted downweighting of correlated or less informative features.

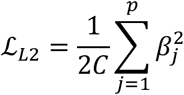

**Equation 1: L2 regularization penalty for logistic regression.** The penalty is calculated from the squared coefficient magnitudes (*β*), where *p* is the number of structural descriptors and *C* is the inverse regularization strength.

Both approaches used class-weighted, L2-regularized logistic regression implemented in scikit-learn (version 1.6.1).^39^ Models were fitted using the lbfgs solver, balanced class weights, a maximum of 5,000 iterations, and a regularization parameter C selected from {0.03, 0.1, 0.3, 1.0}. The model random seed was 42; the solver used its default convergence tolerance. The same algorithm, hyperparameter grid, and inner-validation scoring were used for both approaches. The saved L2 validation audit reported no convergence warnings across its repeated-split and randomized-label fits.

### 3.5 Nested Cross-Validation and Performance Evaluation

Classification was evaluated separately for the four prediction pipelines and three transporter isoforms (12 tasks). Each task used five outer folds for held-out evaluation and five inner folds for feature and hyperparameter selection. Cross-validation partitions were generated using stratified, ligand-grouped splitting with a fixed random seed.

Compound identities were grouped across all three OATP isoforms so that the same compound could not appear in both training and held-out partitions under different OATP labels. The partitions were generated using the complete compound-isoform dataset and shared across all four structure-prediction pipelines, OATP isoforms, and classification approaches, ensuring that performance comparisons were conducted using identical held-out compounds.

Within each outer training partition, candidate model configurations were evaluated using inner cross-validation, with average precision serving as the model-selection criterion. For the individual-metric approach, this procedure selected both the structural metric and classifier hyperparameters that maximized inner out-of-fold average precision. For the combined-metric approach, all eligible structural metrics were supplied as candidate features, and regularization parameters were selected through the same inner-validation framework. Feature eligibility, missing-value imputation, and standardization were determined using the appropriate training partition at each stage of model fitting.

The selected classifier was refitted on the full outer training set and used to predict inhibitor probabilities for its held-out compounds. Concatenating predictions from the five outer folds yielded one out-of-fold probability for every compound within each task and classification approach.

Average precision (AP) and ROC-AUC were used as the primary performance measures. AP is particularly informative when inhibitor and non-inhibitor classes are imbalanced, while ROC-AUC measures discrimination between positive and negative instances across classification thresholds.^40–44^ Additional classification metrics, including balanced accuracy, Matthews correlation coefficient, sensitivity, and precision, were calculated using a probability threshold of 0.5. The primary Top-1 model-integration comparison was the paired difference between the combined-metric and individual-metric classifiers within each predictor-isoform task.

### 3.6 Statistical Comparison of Classification Performance

Differences between individual- and combined-metric classifiers were evaluated using paired statistical procedures applied to their outer-fold predictions. Performance differences were calculated separately for each of the 12 pipeline-isoform tasks and for both primary endpoints.

Uncertainty in Top-1 combined-versus-individual performance differences was estimated using 10,000 paired bootstrap resamples of compound-level held-out predictions. Each resample maintained the correspondence between the predictions from the two classifiers for a compound. Two-sided paired prediction-exchange permutation tests (10,000 exchanges) were used to evaluate task-level differences. The reported 95% confidence intervals are unadjusted and should not, on their own, be interpreted as establishing statistical significance after multiple testing. Adjusted inference is identified explicitly in the Results and figure legends.

To evaluate the overall benchmarking hypothesis, a global effect was calculated as the equally weighted mean performance difference across all 12 pipeline-isoform tasks. Global bootstrap resampling used the same compound-level resample across tasks, thereby preserving the dependence arising from compounds represented in multiple transporter datasets.^45–47^ Global paired permutation tests similarly applied a shared compound-level exchange assignment across tasks. The two global endpoint comparisons were corrected using the Holm procedure.^48^

These inference procedures evaluated uncertainty conditional on the completed nested-cross-validation predictions; model selection and training were not repeated within each bootstrap or permutation iteration.

### 3.7 Robustness and Sensitivity Analysis

#### Cross-validation partition stability

To assess whether the primary Top-1 findings were dependent on one compound allocation, we repeated the complete individual- and combined-metric L2 nested-cross-validation workflow across five fold assignments, including the original assignment. We evaluated the absolute held-out performance of the classifiers as well as the direction and magnitude of their paired differences. These repeated partitions reused the same compounds and therefore provide a stability analysis rather than independent external validation.

#### Randomized-label negative control

To determine whether the modeling and metric-selection procedures could produce apparent classification improvements in the absence of genuine relationships between compound identity and inhibition, 100 randomized-label datasets were constructed. Each randomization reassigned the complete three-transporter inhibition-label profile among compounds, preserving transporter-specific class frequencies while disrupting the association between each compound’s predicted structural features and its inhibition outcomes. The same reassignment was used across all four structure-prediction pipelines, and the definitive top-ranked-structure nested-cross-validation procedure was repeated for each randomized dataset. Performance and combined-minus-individual differences from the experimental labels were compared against the resulting null distributions.

#### Structural-ensemble sensitivity

We evaluated whether incorporating additional predicted complexes improved inhibitor classification beyond the model’s native Top-1 complex. For each of the 12 predictor-isoform tasks, ensemble depth was varied from 1 through 100 structures per compound-isoform pair. The highest-ranked prediction from each of 20 seed groups was selected first, followed by the second-ranked prediction from each seed, etc. This grouping procedure was intended to distribute selection across prediction groups, not to establish that the retained structures were conformationally distinct.

Within each ensemble, a shared regularized logistic model scored the K selected structures, and their predicted probabilities were averaged to yield one probability for the compound-isoform observation. Model fitting optimized a bag-level classification objective, with regularization parameter C selected using AP from five inner held-out folds; each depth was refitted and evaluated on five outer held-out folds. The effect of using ROC-AUC rather than AP as the inner cross-validation optimization criterion was also tested (Supplementary Figure S5). The full depth sweep was repeated across 10 compound-grouped cross-validation partitions using the same data and matched Top-1 baseline within each repeat. For each ensemble size, AP and ROC-AUC differences were computed against Top-1 within the same repeat and then averaged across repeats. The curves are descriptive analyses of held-out performance: the repeated partitions are not independent external cohorts, and peaks selected retrospectively from the depth curves should not be interpreted as validated optimal ensemble sizes.

The Top-1 baseline refers to the native highest-ranked structure from the full set of 100 generated complexes, not to a separate experiment generating only one structure. The ensemble-depth analysis therefore tests the value of incorporating additional existing predictions and does not directly measure savings in structure-generation cost. Because all ensemble depths are evaluated on the same compound collection, apparent changes between neighboring depths must also be interpreted with this repeated-measures dependence in mind.

### 3.8 Descriptor-Family Classification Analysis

To investigate which categories of structural information independently contributed to OATP inhibitor classification, the Top-1 candidate descriptors were assigned to five nonoverlapping groups based on measured properties: interface confidence (ipTM or interface pLDDT), fold/local confidence (pTM or pLDDT), predicted error (PAE or PDE), native ranking/aggregate scores, and clash/disorder indicators. Metric families absent from a given pipeline were not evaluated.

For each of the 12 pipeline-isoform combinations, classifiers were trained using only the structural metrics belonging to one descriptor family at a time, excluding all other candidate features. Each family-only classifier was independently fitted using the same compound-grouped nested cross-validation partitions, training-only preprocessing, and L2-regularized logistic regression framework as the full combined-metric classifier. The regularization parameter was retuned for each family using average precision (AP) within the inner cross-validation folds. Held-out AP and ROC-AUC were calculated for each family-only classifier and compared against the corresponding full combined-metric model. This approach allowed us to evaluate how much predictive performance could be recovered using each descriptor family independently. In a complementary ablation analysis, each descriptor family was excluded from the full combined-metric classifier, with model fitting and hyperparameter selection repeated within the nested cross-validation framework (Supplementary Figure S3).

Uncertainty in absolute family-only performance and paired differences from the full model were estimated using 10,000 compound-level bootstrap resamples of held-out predictions. Two-sided paired prediction-exchange permutation tests were used to evaluate differences between family-only and full-model performance. Benjamini-Hochberg correction was applied within each prediction pipeline and performance endpoint across all available family-isoform comparisons (15 for AlphaFold 3; 12 each for Boltz-2, Chai-1, and Protenix).^43,45^ A broader correction across all 51 available family-only comparisons per endpoint was retained as a sensitivity analysis.

## Results

### 4.1 Benchmark Construction and Structural-Metric Coverage

To systematically evaluate the utility of model-derived structural information for OATP inhibitor classification, we established a common benchmark comprising 224 compounds with experimentally measured inhibition data for OATP1B1, OATP1B3, and OATP2B1. After applying a threshold of 50% inhibition, 170 of the 672 compound-isoform pairs (25.3%) were classified as inhibitors. Inhibitor prevalence varied substantially across transporters, with 78 inhibitors identified for OATP1B1 and 46 each for OATP1B3 and OATP2B1. These differences established transporter-specific classification tasks with distinct positive-class frequencies, an important consideration when evaluating precision-recall performance and comparing the predictive utility of structural descriptors across isoforms. Fingerprint-based chemical similarity analysis indicated broad structural diversity across the 224-compound benchmark, with a median pairwise Tanimoto similarity of 0.09 (Supplementary Figure S6).^49^

For every compound–isoform observation, 100 predicted protein–ligand complexes were assembled from each of four co-folding pipelines. These predictions provided the native highest-ranked structure for the primary single-versus combined-metric benchmark and, separately, the ensembles used to study structural sampling depth. The available candidate descriptors differed among pipelines: 39 for AlphaFold 3, 15 for Boltz-2, 31 for Chai-1, and 74 for Protenix after excluding administrative metadata (Supplementary Table S2). These metrics describe local and global confidence, interfaces, predicted errors, ranking, and steric clashes or disorder. This benchmark therefore tests whether metrics originally intended to describe predicted structural quality also contain information useful for classifying experimental transport inhibition, either alone or in combination. Together, this benchmark establishes a systematic comparison of the functional predictive utility of structural modeling outputs across three pharmacologically relevant OATP isoforms.^30,33–36^

### 4.2 Structural Metrics Predict OATP Inhibition with Task-Dependent Benefits from Integration

Structural co-folding metrics exhibited substantial discrimination of OATP inhibitors from non-inhibitors (Figure 2), even though these descriptors were not developed as functional inhibition predictors. The clearest example was OATP1B3: the optimized single-metric classifier using Chai-1 reached a ROC-AUC of 0.888, approaching the combined-metric performance of 0.893. Across the four prediction pipelines, combined-metric OATP1B3 ROC-AUC values ranged from 0.865 to 0.893, whereas OATP2B1 generally showed lower discrimination (0.701–0.735). For Chai-1/OATP1B3, combined-metric AP was 0.631, appreciably above the 0.205 inhibitor-prevalence baseline. Thus, multiple metrics are not invariably required, but multiple individual outputs of structure-prediction models can carry predictive signal for functional inhibition.

**Figure 1.**
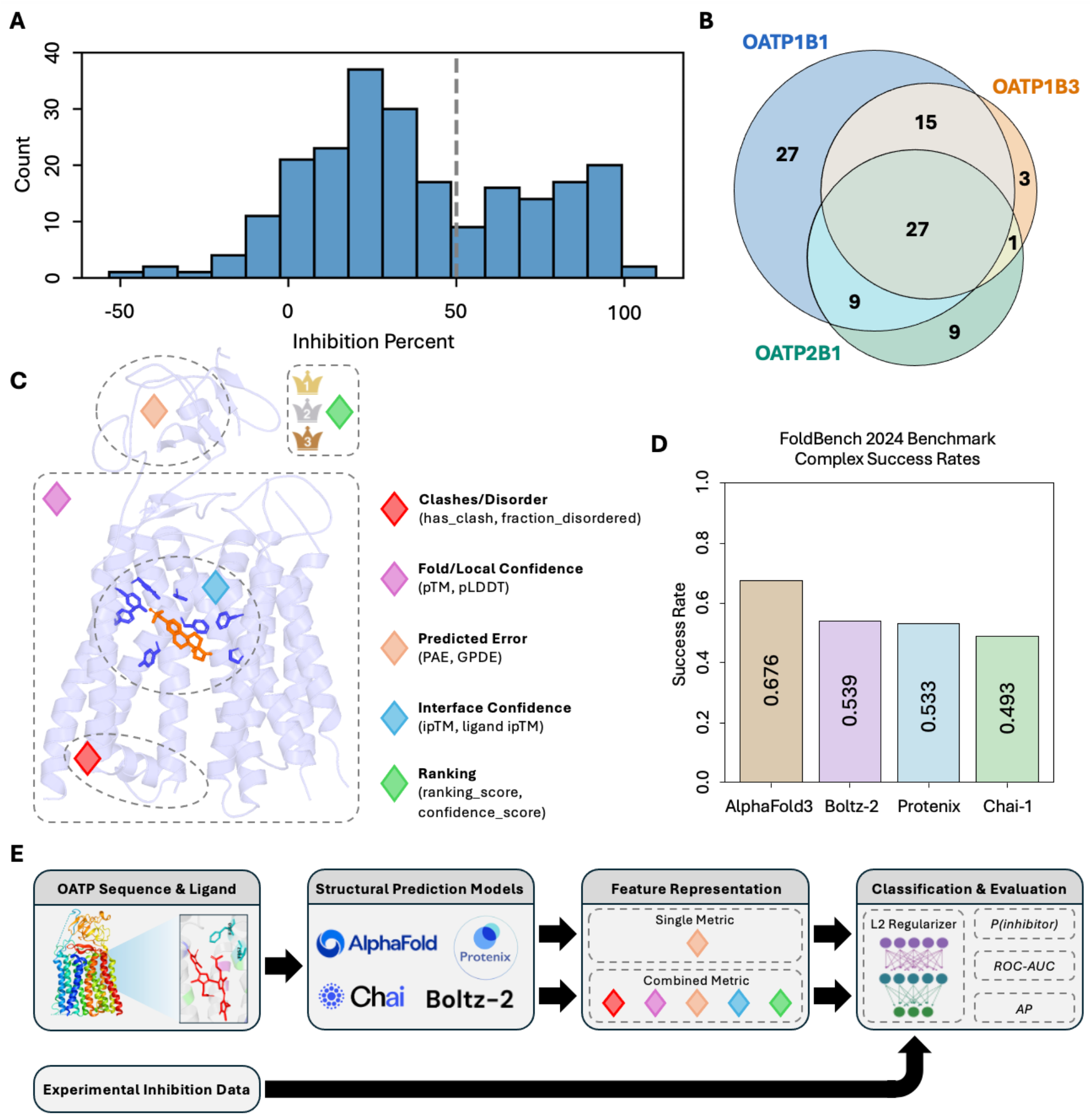
Experimental OATP inhibition benchmark, structural descriptor families, and computational classification workflow. **(A)** Distribution of experimentally measured OATP1B1 inhibition percentages. The dashed vertical line denotes the ≥50% inhibition threshold used to classify compounds as inhibitors. Data from Karlgren et al. (2012).^17^ **(B)** Three-way Venn diagram illustrating inhibitor overlap among OATP1B1, OATP1B3, and OATP2B1. The benchmark comprised 224 compounds evaluated against all three transporters, yielding 672 compound-isoform observations. At the ≥50% inhibition threshold, 78 compounds inhibited OATP1B1, 46 inhibited OATP1B3, and 46 inhibited OATP2B1. Of the 224 compounds, 133 did not inhibit any transporter. **(C)** Experimentally determined OATP1B1 structure in complex with estrone-3-sulfate (E3S; PDB 8HND), illustrating five categories of descriptors extracted from predicted protein-ligand complexes: clashes/disorder, fold/local confidence, predicted error, interface confidence, and ranking/aggregate confidence.^14,15,50^ Colored markers schematically indicate structural regions or properties associated with each descriptor family and do not represent confidence values measured from the experimental structure. TM regions 8 and 10 were omitted for visual clarity. **(D)** Published protein-ligand complex prediction success rates for AlphaFold 3, Boltz-2, Protenix, and Chai-1 from the FoldBench benchmark, using targets released after January 2024. Successful predictions required ligand RMSD <2 Å and protein–ligand interface LDDT >0.8.^51^ These external structural benchmarks are distinct from the OATP inhibitor-classification performance evaluated in this study. **(E)** Overview of the computational inhibitor-classification workflow. OATP protein sequences and candidate ligands are input to four protein-ligand structure-prediction models (AlphaFold 3, Boltz-2, Protenix, and Chai-1). Structural descriptors extracted from the resulting predictions are evaluated individually or combined into multivariable feature representations. Experimental inhibition measurements provide binary classification labels for L2-regularized logistic regression models, which are trained and evaluated using compound-grouped nested cross-validation. Held-out predictions are assessed using average precision (AP) and area under the receiver operating characteristic curve (ROC-AUC).

**Figure 2.**
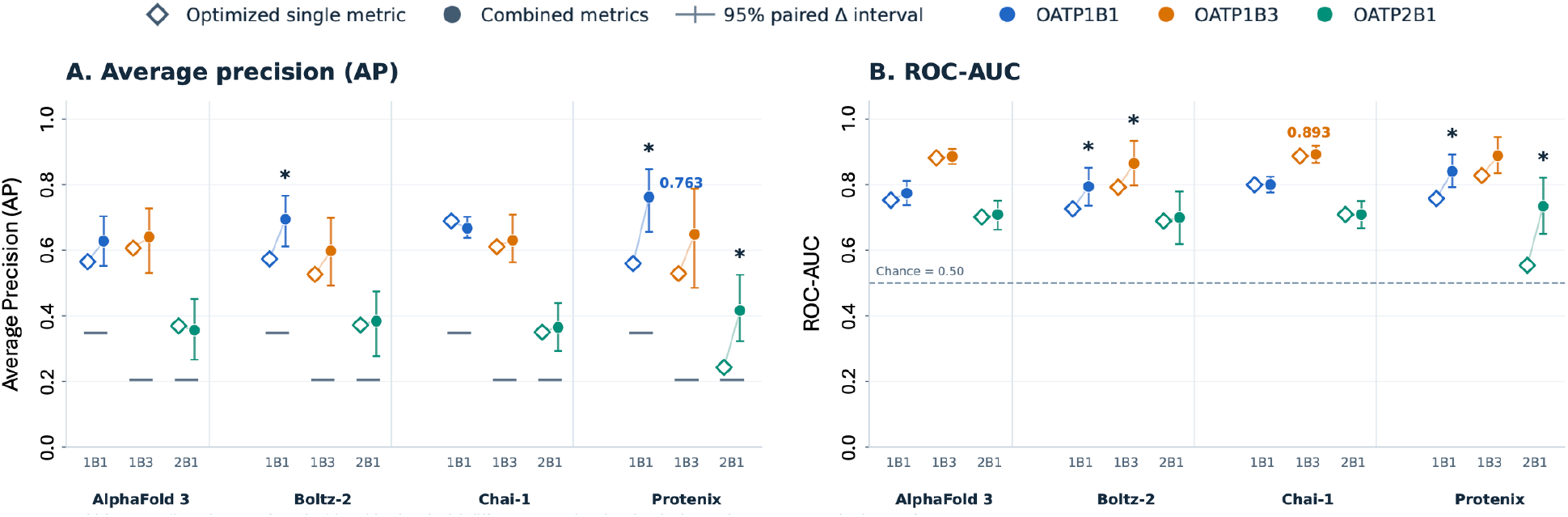
Structural co-folding metrics predict OATP inhibition, with task-dependent additional gains from combining metrics. Held-out **(A)** average precision (AP) and **(B)** ROC-AUC achieved by optimized individual-metric classifiers (open diamonds) and combined-metric classifiers (filled circles) across four protein-ligand structure-prediction pipelines and three OATP isoforms. Colors indicate OATP1B1 (blue), OATP1B3 (orange), and OATP2B1 (teal). Horizontal reference markers in panel A indicate the expected AP under random ranking based on inhibitor prevalence (0.348 for OATP1B1; 0.205 for OATP1B3 and OATP2B1), while the dashed line in panel B indicates chance-level discrimination (ROC-AUC = 0.5). Error bars represent unadjusted 95% paired bootstrap confidence intervals for the performance difference between combined- and individual-metric classifiers. Asterisks indicate statistically significant differences (q < 0.05), determined using two-sided paired permutation tests with Benjamini-Hochberg correction within each model-isoform combination across AP and ROC-AUC. All classifiers were evaluated using the highest-ranked predicted protein-ligand complex (Top-1), a ≥50% experimental inhibition threshold, and five-fold compound-grouped nested cross-validation, with metric selection and hyperparameter optimization performed within training folds. The structural descriptors identified as best-performing for each prediction pipeline and OATP isoform are summarized in Supplementary Table S3.

We next asked whether the numerical signal contained in individual descriptors could be strengthened by adding complementary structural metrics. Effect sizes varied appreciably by predictor and transporter, and only a subset of task-level comparisons met the significance criteria (Figure 2). Protenix showed the largest task-specific benefits from combining metrics. For OATP1B1, AP rose from 0.560 with its optimized individual descriptor to 0.763 with combined metrics (95% CI: 0.656 to 0.848), while ROC-AUC increased from 0.758 to 0.841. For OATP2B1, ROC-AUC increased from 0.554 to 0.735 (95% CI: 0.650 to 0.821), alongside an AP increase from 0.243 to 0.417. Protenix also improved OATP1B3 AP from 0.529 to 0.649, with ROC-AUC changing from 0.828 to 0.889. These results show that metric integration can be particularly helpful for certain pipelines, even though the best individual metrics already yield useful classification performance. Boltz-2 also showed numerical gains from metric integration in selected tasks. For OATP1B1, AP increased from 0.574 to 0.695 and ROC-AUC from 0.727 to 0.795; OATP1B3 ROC-AUC increased from 0.793 to 0.865. In contrast, AlphaFold 3 and Chai-1 generally showed smaller additional benefits, and some tasks had little or no numerical improvement. We therefore separate the finding that structural metrics are informative from the narrower conclusion that integrating metrics improves a specific fitted classifier.

Collectively, these results establish functional predictive value in individual co-folding descriptors and show that additional metrics can sometimes improve that value. The highest absolute discrimination was achieved by Chai-1 and OATP1B3 (ROC-AUC = 0.893), whereas Protenix and OATP1B1 attained the highest combined-metric AP (0.763) and a comparatively large integration benefit. These two findings are complementary rather than competing: a useful single descriptor is already an important result, and a multivariable model offers a task-dependent opportunity for further improvement.

### 4.3 Predictive Robustness Across Cross-Validation Partitions and Randomized Controls

For the Top-1 classification results, we next examined whether performance persisted when compounds were reassigned to different cross-validation folds. First, five repeated compound-grouped nested-CV partitions showed that several strong absolute classifiers remained informative under alternative fold assignments. In particular, Chai-1/OATP1B3 combined-metric ROC-AUC remained between 0.879 and 0.893 across the five folds. Although the combined-metric models generally outperformed the single-metric models, individual pipeline-isoform tasks differed in the size and direction of their improvements. Protenix exhibited improved combined-metric performance in both AP and ROC-AUC for all three OATPs over the five partitions, consistent with the integration benefits observed in the main comparison. For Protenix/OATP1B1, the AP advantage of the combined-metric classifier ranged from +0.143 to +0.255 across partitions; Boltz-2/OATP1B1 also retained positive changes in both endpoints. Other tasks showed less consistent gains, even when their absolute discrimination remained useful. Because the same 224 compounds per isoform were used in every partition, these observations support stability to the split assignment rather than independent validation in new chemistry.

Second, randomized-label controls evaluated whether the observed classification performance depended on the true correspondence between structural features and experimental inhibition. We randomly assigned inhibition label profiles among compounds 100 times, preserving transporter-specific prevalence while disrupting structure-label associations. Across tasks, the descriptive mean combined-metric performance under randomized labels dropped to chance values of 0.264 AP and 0.491 ROC-AUC, compared with 0.564 AP and 0.798 ROC-AUC using experimental labels (empirical p = 0.0099 for both aggregate performance comparisons). These aggregate figures summarize the negative control, not uniform performance across individual transporters or pipelines. The observed performance improvements, null distributions, and stability across repeated cross-validation partitions are summarized in Supplementary Figure S7.

Together, the repeated-partition and randomized-label analyses address different aspects of robustness: the former indicates whether the observed predictive relationships are sensitive to fold assignment, whereas the latter shows that the signal is completely diminished when experimental inhibition labels are decoupled from structural descriptors. Neither analysis constitutes independent external validation of the data. Nonetheless, the controls support investigating how different representations of the predicted structural ensembles affect classification.

### 4.4 Structural Ensemble Depth Produces Pipeline-Dependent Effects

For each compound-isoform pair, we generated 100 complexes across each model, allowing us to ask whether additional predictions improved inhibitor classification relative to using only the highest-ranked complex. For each predictor and isoform, we fitted compound-level bag classifiers over every ensemble depth from 1 to 100 structures and evaluated held-out AP and ROC-AUC over 10 matched, repeated compound-grouped nested-CV partitions (Figure 3). Each model made predictions across 20 seeds for each predictor-isoform pair, with 5 poses each. We used tiered ranking to add the highest-ranked pose from each seed to the classifier first before progressively adding lower-ranked poses from each seed. These 100 predictions may contain redundant or related structures, and our grouping does not establish conformational diversity.

**Figure 3.**
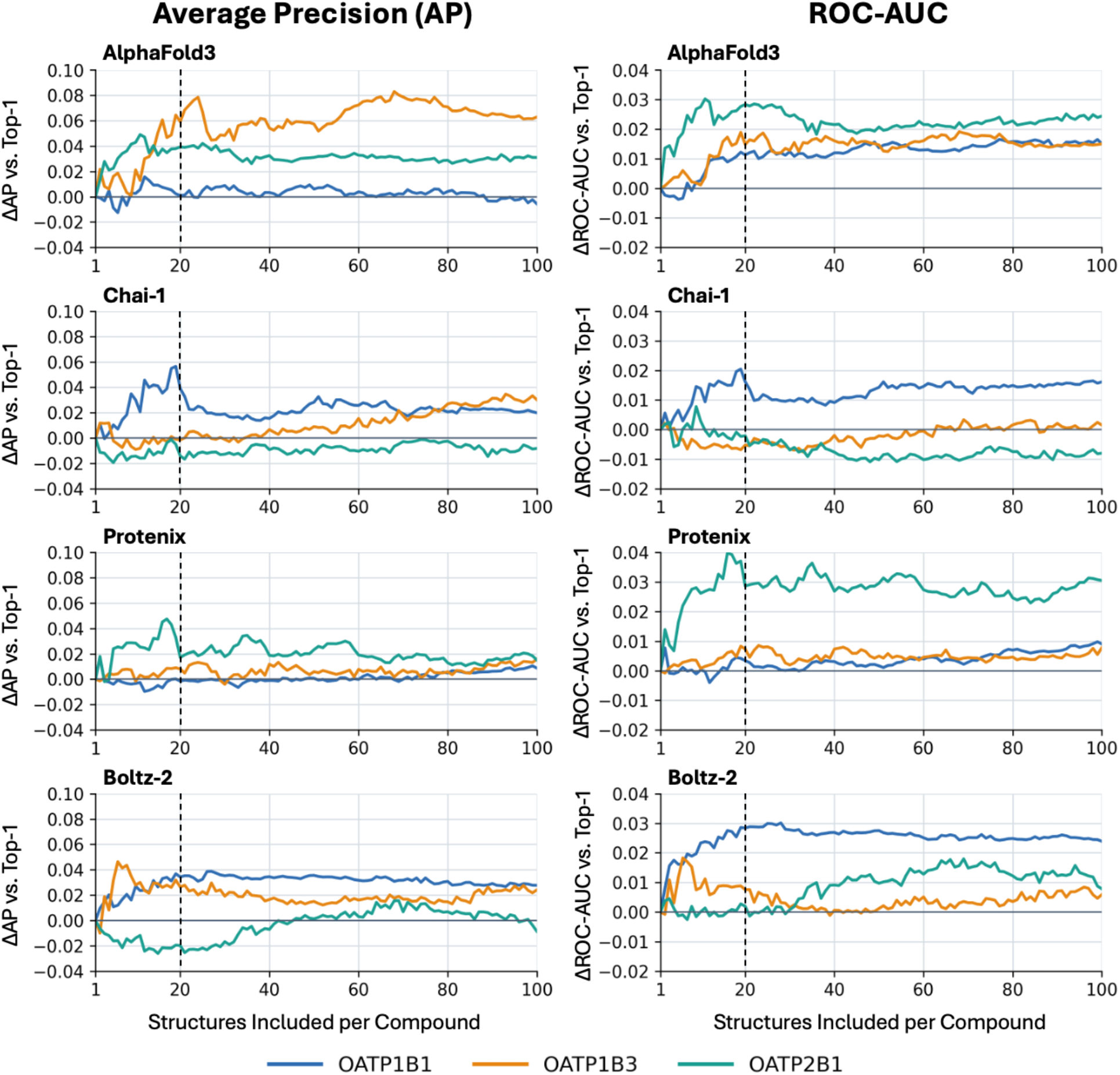
Structural ensemble sampling produces model-dependent effects on OATP inhibitor classification. Changes in held-out average precision (left) and ROC-AUC (right) as increasing numbers of predicted protein-ligand complexes are incorporated into the classifier. Results are shown for AlphaFold-3, Chai-1, Protenix, and Boltz-2 (top to bottom), across OATP1B1 (blue), OATP1B3 (orange) and OATP2B1 (teal). For each model, structures were incorporated using a tiered ranking strategy, prioritizing the highest-confidence prediction from each of 20 seeded groups before incorporating progressively lower-ranked predictions from those groups. Performance was evaluated across ensemble depths of 1-100 structures per compound using repeated, compound-grouped nested cross-validation. Curves represent the mean change in classification performance across 10 cross-validation repetitions relative to classifiers trained using only the Top-1 ranked structure. Positive values indicate improved classification performance relative to Top-1.

The contribution of additional structures varied substantially by predictor, isoform, and performance endpoint. Some curves improved early and then changed relatively little with further poses, while others continued to improve or declined after intermediate depths. For AF3/OATP1B3, AP improved by +0.059 relative to Top-1 at 20 poses and +0.063 at 100 poses; the corresponding ROC-AUC improvements were approximately +0.015 at both depths. AF3/OATP2B1 likewise showed positive changes at 20 poses (ΔAP = +0.039; ΔROC-AUC = +0.028), with smaller AP gains at 100 poses (+0.031). Thus, some of the additional discriminatory information was accessible without incorporating all 100 predictions, but the depth-response profiles were not uniformly monotonic.

Several informative patterns appeared across individual tasks. Boltz-2/OATP1B1 improved by ΔAP = +0.034 and ΔROC-AUC = +0.028 at 20 structures, with slightly smaller improvements at 100 structures (+0.028 AP and +0.024 ROC-AUC). Chai-1/OATP1B1 showed a relatively early AP benefit at 20 structures (+0.039 AP) that diminished at 100 structures (+0.020 AP), even though its ROC-AUC gain remained near +0.016 at both depths. For Protenix/OATP2B1, ROC-AUC increased by +0.029 at 20 structures and +0.031 at 100 structures, while AP improvements were more modest (+0.017 AP and +0.015 AP, respectively). The differing behavior of AP and ROC-AUC indicates that additional structures can alter high-ranked inhibitor retrieval without producing proportional changes in global discrimination, or vice versa.

Other combinations showed little benefit or worsened with greater ensemble depth. Chai-1/OATP2B1 showed minimal losses in ΔAP and ΔROC-AUC at 100 structures (both −0.008), whereas Chai-1/OATP1B3 showed a larger ΔAP gain at 100 structures (+0.030 AP) than at 20 structures (−0.001 AP), despite a comparatively small change in ROC-AUC at 100 structures (+0.001). Protenix/OATP1B1 showed minimal ΔAP loss at 20 structures (−0.002 AP) and gained only +0.009 AP at 100 structures. Such differences caution against assuming a universally optimal ensemble depth or interpreting more predictions as necessarily more functional information.

Several predictor-isoform combinations began to approach stable performance around 20 structures, right at the point at which the highest-ranked prediction from each seed or index group had been incorporated. This observation suggests that the initial predictions from different groups may capture much of the structural information relevant to inhibitor classification, while additional predictions from previously represented groups provide diminishing benefits. Similar observations have been reported in protein-complex structure prediction, where increasing the number of independent prediction seeds improved structural accuracy, while generating additional diffusion samples within individual seeds provided comparatively limited benefits.^24^ More recently, it has been demonstrated that sampling across different seeds produced greater structural diversity than generating additional predictions within the same seed.^52^ Although these findings provide a possible explanation for the diminishing returns observed here, our analysis did not directly evaluate structural similarity or independence among predicted complexes. Moreover, the plateau was not universal, with some predictor-isoform combinations continuing to improve beyond 20 structures. Thus, these results suggest that sampling across seed groups may provide complementary predictive information, but do not establish that additional structures are redundant or that 20 predictions represent an optimal ensemble size.

Overall, ensemble-depth effects were task-specific and frequently exhibited diminishing returns before all 100 predicted structures had been included. This plateau is consistent with, but does not demonstrate, redundancy among predicted complexes; it may also reflect uneven prediction quality or limitations of equal-weight probability aggregation. We did not directly quantify structural independence in this analysis, and the curves are descriptive summaries of repeated evaluations on the same compound collection, not formal evidence that a particular number of structures is optimal. The results instead indicate that both the choice of structure-ordering strategy and the way individual predictions are aggregated warrant careful evaluation when applying co-folding outputs to functional classification.

### 4.5 Predictive Utility of Structural Descriptor Families Varies Across Modeling Pipelines

To investigate which categories of structural information were most informative for OATP inhibitor classification, we evaluated whether individual descriptor families could independently predict inhibition. Structural metrics were grouped into five categories: predicted error, fold/local confidence, interface confidence, ranking/aggregate scores, and clash/disorder indicators. For each prediction pipeline and OATP isoform, classifiers were trained using only the metrics within a single family. These family-only classifiers were independently optimized using the same nested cross-validation framework and compared against the full combined-metric classifiers (Figure 4). This approach allowed us to evaluate which types of structural information were sufficient to support inhibitor classification without requiring integration of all available metrics.

**Figure 4.**
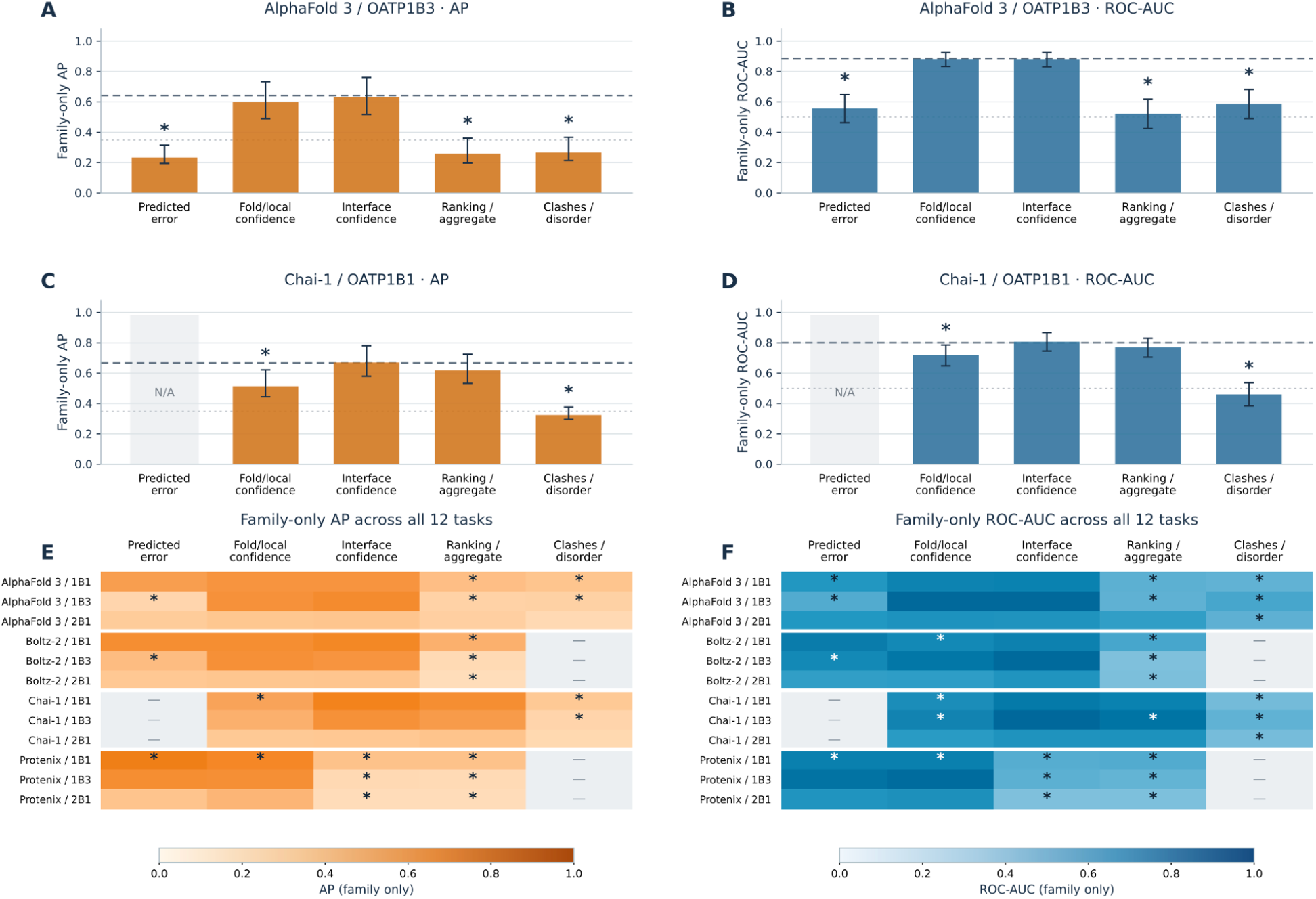
Individual structural descriptor families exhibit varying predictive utility for OATP inhibitor classification. (A-D) Held-out average precision (AP) and ROC-AUC achieved by classifiers trained using only one structural descriptor family at a time for AlphaFold 3/OATP1B3 **(A, B)** and Chai-1/OATP1B1 **(C, D)**. Each classifier was independently trained and optimized using only the descriptors belonging to the indicated family, excluding all other structural metrics. Bars represent absolute classification performance, with error bars indicating unadjusted 95% confidence intervals from 10,000 compound-level bootstrap resamples. Dashed horizontal lines indicate the performance of the full combined-metric classifier, while dotted lines indicate random-ranking baselines (AP equal to inhibitor prevalence; ROC-AUC = 0.5). **(E, F)** Heatmaps summarize family-only AP **(E)** and ROC-AUC **(F)** across all 12 model-isoform combinations. Color intensity indicates absolute classification performance, while gray cells represent descriptor families unavailable for a given prediction pipeline. Asterisks indicate statistically significant differences between family-only and full combined-metric classifiers (q < 0.05), determined using two-sided paired permutation tests with 10,000 prediction exchanges and Benjamini-Hochberg correction within each prediction pipeline and performance endpoint across all available descriptor-family comparisons. All analyses used the highest-ranked predicted protein-ligand complexes (Top-1) and compound-grouped nested cross-validation, with feature selection and hyperparameter optimization restricted to training folds. Complementary leave-one-family-out analyses further evaluated the effects of excluding individual descriptor families from the full combined-metric classifier (Supplementary Figure S3).

Several descriptor families independently achieved strong classification performance, particularly for OATP1B3. For AlphaFold3/OATP1B3, interface-confidence metrics alone achieved an average precision (AP) of 0.632 and ROC-AUC of 0.882, mirroring the full combined-metric classifier (AP = 0.641; ROC-AUC = 0.886). Fold/local-confidence metrics produced similarly strong results (AP = 0.600; ROC-AUC = 0.882), with neither family showing a statistically significant difference from the full model. In contrast, predicted-error, ranking/aggregate, and clash/disorder features performed substantially worse in isolation, with ROC-AUC values ranging from 0.521 to 0.588. These differences were statistically significant following multiple-testing correction (q < 0.05), suggesting that the predictive information available from AlphaFold 3 was concentrated primarily within its interface- and fold-confidence descriptors.

Chai-1 exhibited a similar dependence on interface-confidence information. For OATP1B1, the interface-confidence-only classifier achieved AP = 0.670 and ROC-AUC = 0.807, closely matching the full combined-metric model (AP = 0.668; ROC-AUC = 0.801). In contrast, fold/local-confidence descriptors alone produced lower performance (AP = 0.514; ROC-AUC = 0.719), with both reductions statistically significant after correction. Interface-confidence metrics were also highly informative for Chai-1/OATP1B3, achieving ROC-AUC = 0.885 compared with 0.893 for the full classifier. Together, these findings indicate that interface-confidence descriptors alone can recover much of the predictive performance achieved by combining all available Chai-1 metrics.

In contrast, Protenix exhibited a greater benefit from integrating information across descriptor families. For OATP1B1, predicted-error metrics were the strongest individual family by AP, achieving 0.688, followed by fold/local-confidence metrics at 0.648. Although both families independently supported inhibitor classification, neither reproduced the full combined-metric performance of 0.763 AP, with statistically significant reductions for predicted-error and fold/local-confidence models. Similar differences were observed for ROC-AUC, where both family-only classifiers achieved values of approximately 0.778 compared with 0.841 for the full model. These results suggest that Protenix benefits from combining information that is not fully captured by either descriptor family independently.

Across all 12 model–isoform combinations, the predictive utility of individual descriptor families varied substantially (Figure 4E,F). Interface- and fold/local-confidence metrics frequently supported strong classification performance, but no single descriptor family consistently reproduced the full classifier across all prediction pipelines and transporters. Importantly, the ability of a family to perform well independently does not establish that it contributes unique information when combined with other descriptors. Likewise, differences in family-only performance should not be interpreted as evidence that particular confidence measures directly capture the molecular mechanisms of transport inhibition. Rather, these findings demonstrate that useful functional information is distributed differently across the outputs of modern co-folding models. For some classification tasks, a relatively limited set of related structural metrics may be sufficient, whereas others benefit from integrating multiple categories of structural information.

## Discussion

Modern protein-ligand co-folding pipelines provide more than candidate structures: their numerical metadata can also serve as features for experimentally grounded functional prediction. Here, we demonstrate that structural metrics contain signal strongly associated with OATP inhibition across four prediction pipelines and three OATP transporters, despite not being explicitly designed for functional predictive purposes. Chai-1/OATP1B3 provides a particularly clear example, with an optimized individual descriptor (ipTM) achieving ROC-AUC = 0.888 and combined-metric performance reaching 0.893. The strong single-metric result is itself useful: a simple confidence-derived feature can be predictive of a functional endpoint without being interpreted as a direct measurement of binding affinity, inhibitory potency, or the transport mechanism.^4,16,24,28^

Combining multiple metrics offered an additional, but not universal, opportunity to improve classification. Protenix/OATP1B1, for example, improved from 0.560 to 0.763 AP after metric integration, while other predictor-isoform combinations showed substantially smaller numerical changes. Repeated cross-validation supported the stability of several strong classifiers to alternative fold assignments, and randomized-label controls showed that the structural-metric performance approached random noise when experimental inhibition labels were decoupled from the predictions. Descriptor-family ablations further suggested that different information categories contribute more across pipelines: error and local-confidence metrics were useful for Protenix/OATP1B1, whereas interface-confidence metrics were particularly informative for Chai-1/OATP1B3. Together, these results demonstrate the value of evaluating both optimized single descriptors and integrated feature sets, rather than assuming that higher model complexity necessarily improves performance.

Using multiple predicted complexes yielded varied benefits. In the ensemble depth analysis, some tasks improved after a relatively small number of structures and then approached a plateau, whereas others continued to change or declined as more predictions were included. The predictor-specific response to ensemble depth suggests that simply generating or using more structures is not a reliable strategy for obtaining better functional classification. At the same time, a performance plateau does not prove structural redundancy: differences in prediction quality, confidence rankings, and pose probabilities could also be limiting gains. These findings motivate future comparisons of conformational diversity and alternative approaches that weight or select structures based on their complementary information content.^42–44,53,54^

This work has several limitations that define its next steps. The benchmark contains 224 compounds per transporter from one experimental study, and even extensive internal compound-grouped cross-validation cannot establish generalization to other inhibitor libraries. The binary endpoint reflects a ≥50% inhibition cutoff at a particular experimental concentration and is not a clinical DDI threshold. Sensitivity to alternative thresholds was evaluated (Supplementary Figure S2), but the classification endpoint remains assay-dependent. Co-folding confidence metrics are indirect descriptors rather than measurements of affinity, transport kinetics, or mechanism. External and prospective experimental validation, comparisons with ligand-only baselines, and evaluation of explicit transporter-ligand interactions will be important tests of generalizability.^4,16,28,36,53^ Ultimately, this study establishes an experimentally grounded framework for evaluating the functional utility of modern protein-ligand structure predictions, highlighting their promise for transporter inhibitor classification and the importance of model-specific evaluation.

## Data Availability

The code, structural prediction metadata, experimental benchmark, and analysis results supporting this study are available at the Woldring Lab GitHub repository: https://github.com/WoldringLabMSU/OATP_Pipeline. The repository includes workflows for protein-ligand complex prediction, OATP inhibitor classification, statistical validation, and manuscript figure reproduction.

