## Supplementary Figures S1-S7 and Tables S1-S3 for "Structural Metrics from Predicted Protein-Ligand Complexes Enable OATP Inhibitor Classification"

**Supplemental Figure S1:** Distribution of experimental OATP inhibition measurements.

**Supplemental Figure S2:** Sensitivity of inhibitor classification to experimental inhibition threshold from 25 to 50%.

**Supplemental Figure S3:** Structural Descriptor Family Ablation from Full Combined-Metric Classifier

**Supplemental Figure S4:** Linear Relationships Between Experimental Inhibition and High-Profile Co-Folding Metrics.

**Supplemental Figure S5:** Sensitivity of OATP inhibitor classification to the inner cross-validation optimization criterion.

**Supplemental Figure S6:** Fingerprint-based chemical similarity across the 224-compound OATP inhibition dataset.

**Supplemental Figure S7:** Permutation controls and repeated cross-validation demonstrate robustness of OATP inhibitor classification improvements.

**Supplemental Table S1:** Inhibition benchmark composition

**Supplemental Table S2:** Structural-metric inventory by model

**Supplemental Table S3:** Structural descriptors selected by optimized single-metric classifiers

### Reproducibility and Code Availability

The computational workflows and supporting datasets used in this study are available at https://github.com/WoldringLabMSU/OATP_Pipeline. The repository contains documentation for the four protein–ligand co-folding pipelines (AlphaFold 3, Boltz-2, Chai-1, and Protenix), structural descriptors, classifier evaluation, statistical analyses, and reproduction of the supplemental figures.

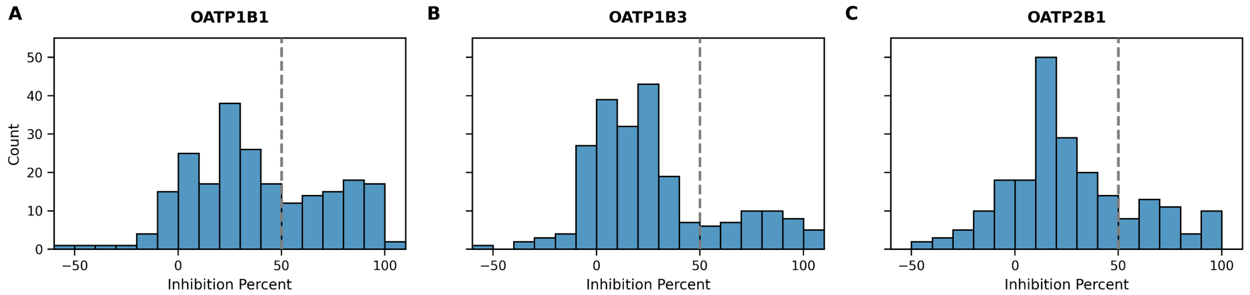


**Supplemental Figure S1: Distribution of experimental OATP inhibition measurements.** Histograms show inhibition percentages for 224 compounds against **(A)** OATP1B1, **(B)** OATP1B3, and **(C)** OATP2B1, using experimental data from Karlgren et al. (2012). Dashed vertical lines indicate the 50% inhibition threshold used for binary inhibitor classification. All panels use identical bin widths and axis scales. For visualization, inhibition values are displayed from −60% to 110%, excluding one OATP1B3 and nine OATP2B1 measurements outside this range.

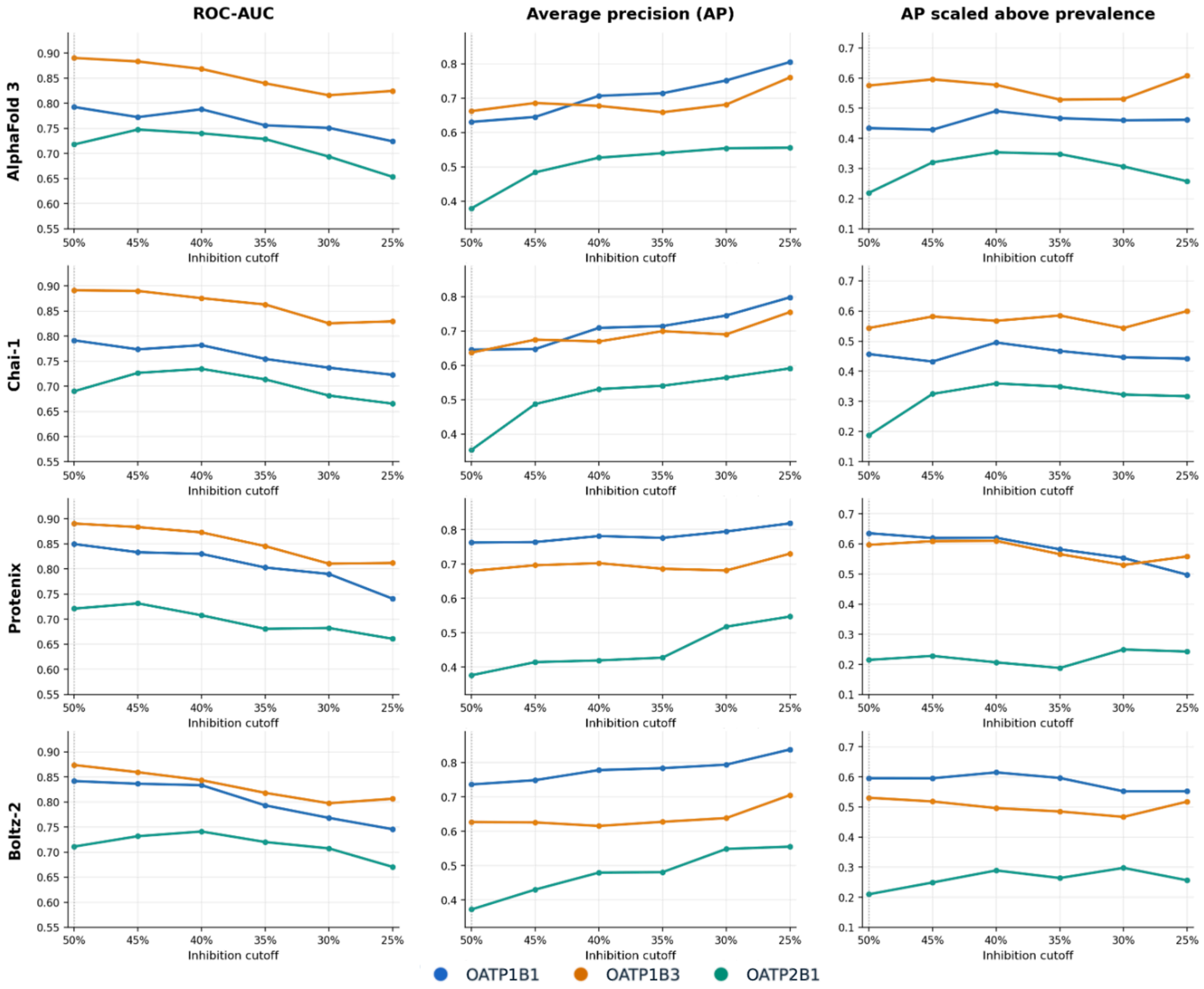


**Supplemental Figure S2: Sensitivity of inhibitor classification to experimental inhibition threshold from 25 to 50%.**

Across four pipelines, each panel displays ROC-AUC, raw AP, or AP scaled above the class-prevalence baselines. Colors denote each isoform, OATP1B1 (blue), OATP1B3 (orange) and OATP2B1 (teal). Curves are descriptive means over 10 repeated compound-grouped nested CV-partitions; the underlying positive-class definition changes across thresholds.

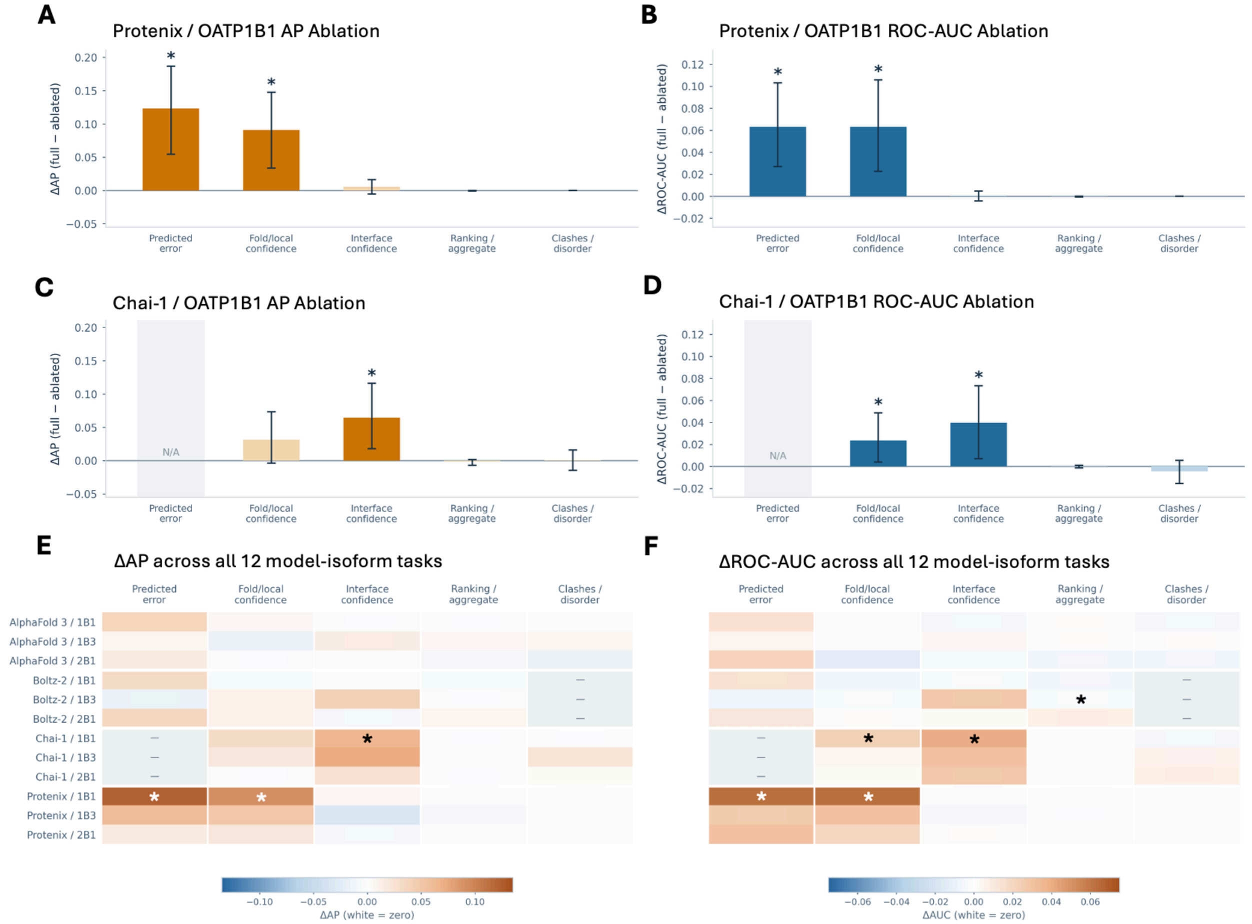


**Supplemental Figure S3: Structural Descriptor Family Ablation from Full Combined-Metric Classifier**

We grouped descriptors into five major categories: predicted interface confidence, fold/local confidence, predicted error, aggregate ranking, and clash/disorder information. Each group was systematically excluded from the combined-metric classifier, with model fitting and hyperparameter selection repeated within the nested cross-validation framework.

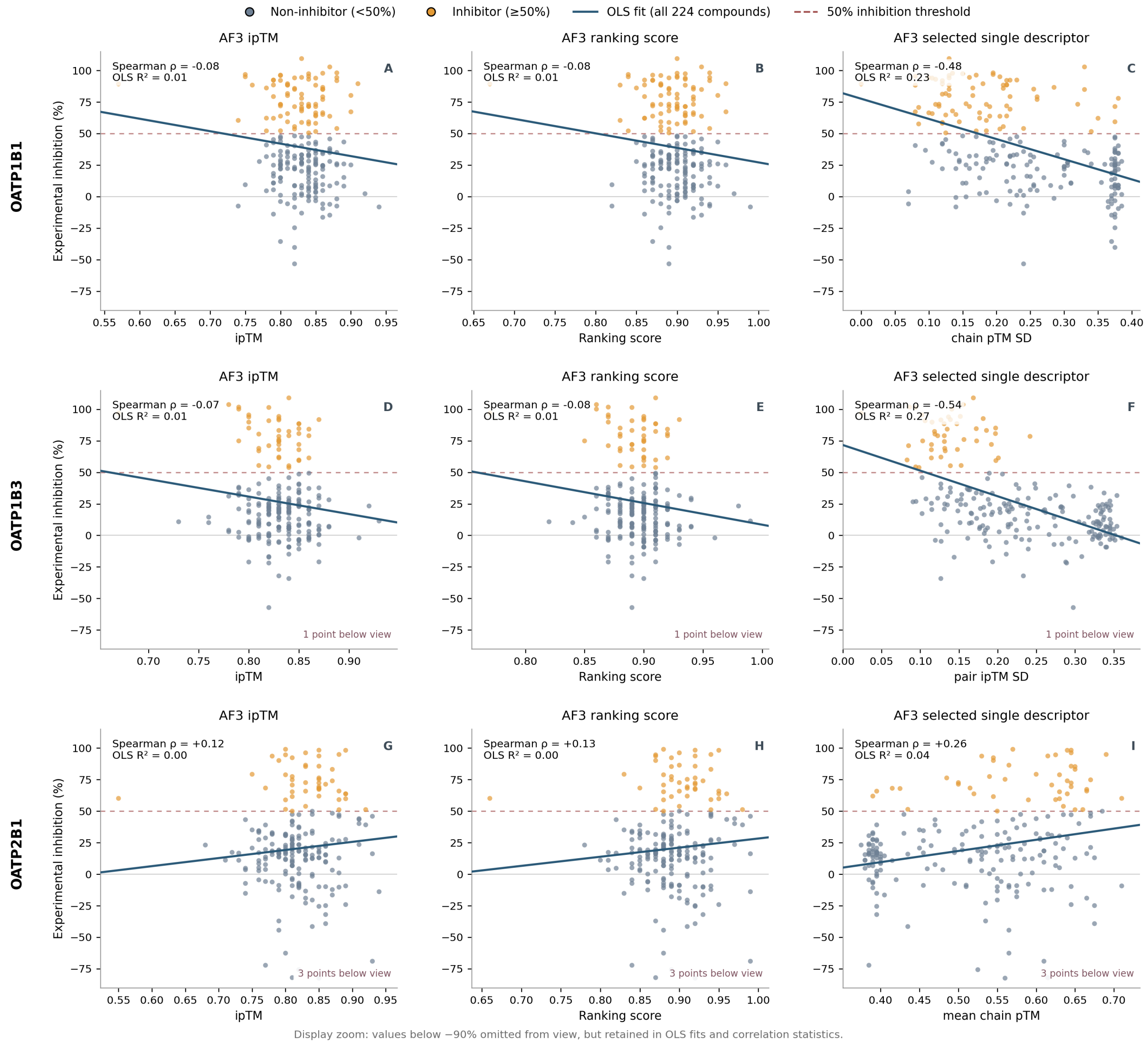


**Supplemental Figure S4: Linear Relationships Between Experimental Inhibition and High-Profile Co-Folding Metrics.**

Plots of Top-1 AlphaFold3 co-folding data from OATP1B1 (top), OATP1B3 (middle), and OATP2B1 (bottom) against ipTM (left), ranking score (middle), and the highest selected single descriptor for each run (right). No meaningful linear relationships were observed.

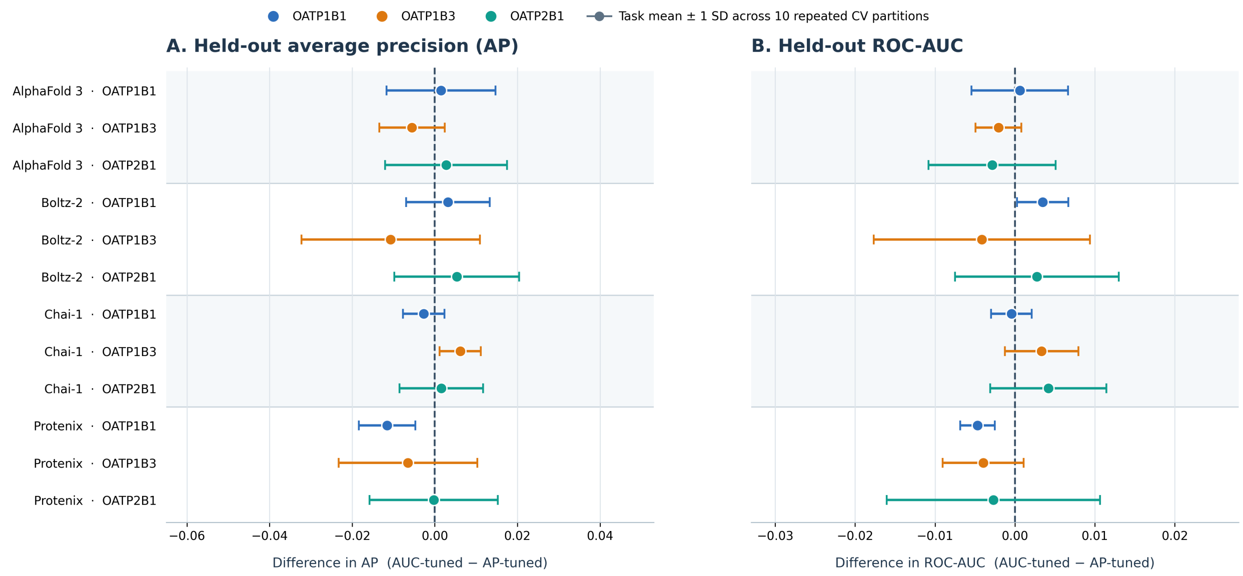


**Supplemental Figure S5: Sensitivity of OATP inhibitor classification to the inner cross-validation optimization criterion.**

Changes in held-out **(A)** average precision (ΔAP) and **(B)** ROC-AUC (ΔROC-AUC) when ROC-AUC rather than AP was used for hyperparameter selection. Results are shown for AlphaFold 3, Boltz-2, Chai-1, and Protenix across OATP1B1 (blue), OATP1B3 (orange), and OATP2B1 (teal), using bag-level classifiers with 100 predicted complexes per compound. Points represent mean differences across 10 repeated compound-grouped nested cross-validation partitions, with error bars indicating standard deviation. Negative values favor AP optimization, while dashed vertical lines indicate no difference. The bottom summary compares Top-1, Top-20, and Top-100 classifiers. Overall, ROC-AUC optimization produced small, task-dependent differences and slightly lower average held-out performance without demonstrating any advantages over AP optimization.

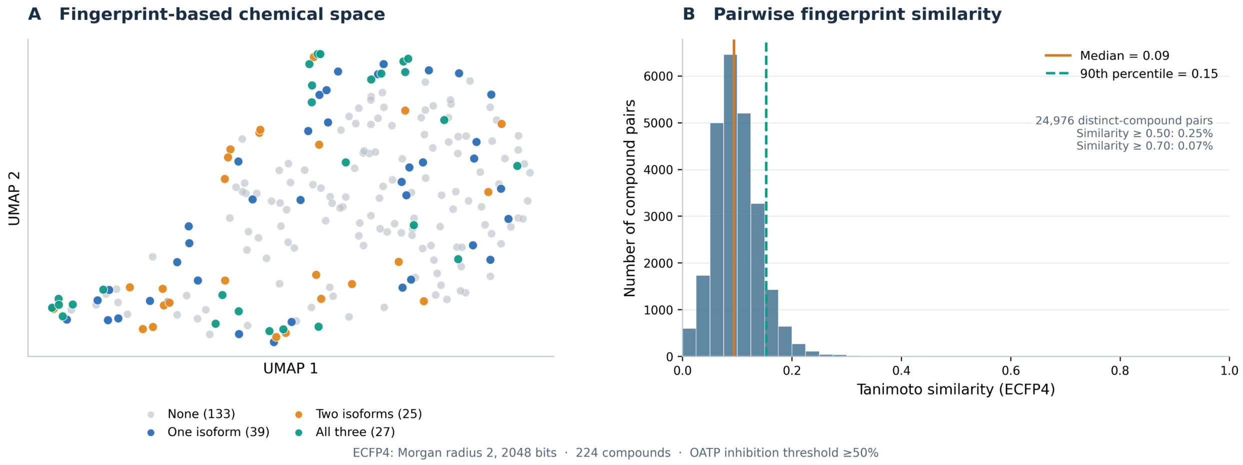


**Supplemental Figure S6: Fingerprint-based chemical similarity across the 224-compound OATP inhibition dataset. (A)** UMAP projection of pairwise ECFP4 Tanimoto distances, with compounds colored by the number of OATP isoforms inhibited at the ≥50% inhibition threshold. **(B)** Distribution of pairwise ECFP4 Tanimoto similarities across all 24,976 unique compound pairs. Vertical lines indicate the median (0.09) and 90th-percentile (0.15) similarities. The low overall pairwise similarity indicates broad chemical diversity within the dataset.

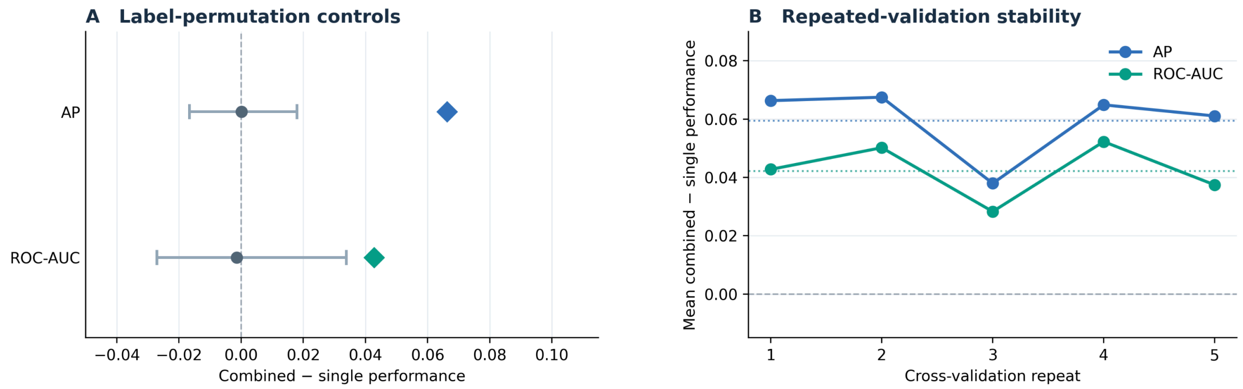


**Supplemental Figure S7: Permutation controls and repeated cross-validation demonstrate robustness of OATP inhibitor classification improvements. (A)** Observed improvements in average precision (AP) and ROC-AUC from combining structural metrics compared with single-metric classifiers, averaged across 12 model–isoform combinations. Colored diamonds indicate observed improvements, while gray circles and error bars represent the mean and central 95% interval of the corresponding null distributions generated from 100 label permutations. **(B)** Mean improvements in AP and ROC-AUC across five repeated nested cross-validation runs. Dotted horizontal lines indicate the mean improvement across repeats. Positive values indicate improved classification performance using combined structural metrics.

**Supplemental Table S1: Inhibition Benchmark Composition**

**A.** Dataset size and class prevalence at the primary ≥50% inhibition threshold.

| Transporter | Observations | Inhibitors | Non-Inhibitors | Prevalence |
|---|---|---|---|---|
| OATP1B1 | 224 | 78 | 146 | 34.8% |
| OATP1B3 | 224 | 46 | 178 | 20.5% |
| OATP2B1 | 224 | 46 | 178 | 20.5% |
| Total | 672 | 170 | 502 | 25.3% |

**B.** Mutually exclusive inhibitor profiles across the same 224 compounds between OATP isoforms.

| Inhibition Profile | Number of Compounds | Percentage of Compounds |
|---|---|---|
| None | 133 | 59.4% |
| OATP1B1 | 27 | 12.1% |
| OATP1B3 | 3 | 1.3% |
| OATP2B1 | 9 | 4.0% |
| OATP1B1 & OATP1B3 | 15 | 6.7% |
| OATP1B1 & OATP2B1 | 9 | 4.0% |
| OATP1B3 & OATP2B1 | 1 | 0.4% |
| All Three | 27 | 12.1% |
| Total | 224 | 100.0% |

**C.** Inhibition % effect on OATP1B1, OATP1B3, and OATP2B1 from Karlgren et al. (2012).

| Compound | OATP1B1 Inhib % | OATP1B3 Inhib % | OATP2B1 Inhib % |
|---|---|---|---|
| Atazanavir | 95.0 | 92.6 | 90.1 |
| Atorvastatin | 96.0 | 74.4 | 98.3 |
| Bromosulfalein | 94.1 | 74.9 | 69.3 |
| Cholecystokinin 8 | 89.3 | 96.9 | 60.1 |
| Dipyridamole | 91.9 | 91.8 | 83.1 |
| Fluo-3 | 93.9 | 88.5 | 65.9 |
| Fluvastatin | 80.1 | 76.0 | 98.2 |
| Glycochenodeoxycholate | 80.2 | 72.0 | 65.3 |
| Glycodeoxycholate | 85.0 | 79.0 | 53.8 |
| Indocyanine green | 86.1 | 100.1 | 79.2 |
| Lopinavir | 85.8 | 84.6 | 86.4 |
| Mifepristone | 81.2 | 80.3 | 70.7 |
| MK-571 | 88.5 | 80.7 | 75.2 |
| Morin | 85.2 | 78.5 | 71.3 |
| Novobiocin | 58.1 | 84.2 | 85.6 |
| Pitavastatin | 97.4 | 94.3 | 63.7 |
| Rifamycin SV | 95.4 | 101.4 | 74.8 |

| | | | |
|---|---|---|---|
| Ritonavir | 92.3 | 85.1 | 93.6 |
| Rosuvastatin | 71.4 | 55.4 | 51.2 |
| Silymarin | 94.5 | 88.5 | 74.0 |
| Sulfasalazine | 92.1 | 72.1 | 92.3 |
| Taurocholate | 93.0 | 95.5 | 96.2 |
| Taurodeoxycholate | 81.7 | 61.4 | 85.3 |
| Taurolithocholate | 97.8 | 89.8 | 94.6 |
| Telmisartan | 109.4 | 91.5 | 94.9 |
| Tipranavir | 89.5 | 109.0 | 99.0 |
| 5-Carboxyfluorescein diacetate | 70.7 | 71.1 | -24.9 |
| Benzbromarone | 86.6 | 19.5 | 76.3 |
| Budesonide | 73.2 | 79.0 | 30.3 |
| Cerivastatin | 73.6 | 68.1 | 40.1 |
| Clarithromycin | 73.1 | 53.8 | 5.1 |
| Cyclosporin A | 96.8 | 103.7 | 14.0 |
| Diazepam | 51.6 | 9.3 | 51.4 |
| Diethylstilbestrol | 62.1 | 31.1 | 68.1 |
| Estradiol-17-β-glucuronide | 67.9 | 69.0 | -12.2 |
| Genistein | 84.9 | -5.0 | 67.9 |
| GF120918 (Elacridar) | 94.4 | 37.4 | 67.2 |
| Glibenclamide | 92.4 | 49.3 | 77.4 |
| Glycyrrhizic acid | 65.8 | 90.9 | 28.7 |
| Ivermectin | 64.7 | 55.2 | 39.0 |
| KO143 | 59.3 | 24.2 | 78.8 |
| Nefazodone | 12.0 | 61.4 | 61.5 |
| Nelfinavir | 71.3 | 59.3 | 50.0 |
| Nystatin | 69.3 | 74.9 | 23.4 |
| Paclitaxel | 71.6 | 62.1 | 30.8 |
| PSC833 (Valspodar) | 96.3 | 93.9 | 33.1 |
| Quercetin | 77.0 | 21.6 | 72.6 |
| Repaglinide | 88.4 | 83.2 | 42.2 |
| Reserpine | 67.2 | 25.4 | 72.3 |
| Rifampicin | 88.3 | 101.7 | 21.2 |
| Taurochenodeoxycholate | 89.5 | 82.4 | 47.2 |
| Vinblastine | 57.6 | 54.3 | -9.3 |
| 17β-estradiol | 102.8 | 39.3 | 47.4 |
| Amprenavir | 79.3 | 16.3 | -3.4 |
| Astemizole | 22.3 | 24.4 | 58.9 |
| Baicalin | 27.6 | 20.9 | 59.1 |
| Candesartan | 52.1 | 28.6 | 28.4 |
| Coumestrol | 73.2 | -21.1 | -41.5 |

| | | | |
|---|---|---|---|
| Diclofenac | 77.9 | 27.3 | 19.6 |
| Erlotinib | 10.6 | 27.5 | 93.7 |
| Erythromycin | 58.8 | 45.8 | -19.0 |
| Estrone-3-sulphate | 98.1 | 10.1 | 20.8 |
| Ezetimibe | 55.5 | 1.5 | -8.9 |
| Flutamide | 6.8 | 9.0 | 65.8 |
| Gemfibrozil | 59.3 | 14.4 | 15.1 |
| Glycocholic acid | 66.4 | 29.0 | -9.8 |
| Hoechst 33342 | 48.1 | 67.0 | 12.7 |
| Indinavir | 71.9 | 18.1 | 17.8 |
| Indometacin | 88.6 | 48.6 | -82.4 |
| Itraconazole | 22.3 | -2.3 | 59.8 |
| Ketoconazole | 53.6 | 24.7 | 44.0 |
| Levothyroxin | -13.1 | 11.4 | 63.7 |
| Lovastatin | 66.5 | 21.3 | -17.9 |
| Mitoxantrone | 27.2 | 67.9 | 48.3 |
| Nicardipine | 65.1 | 10.1 | 31.8 |
| Nifedipine | 63.7 | -21.0 | -44.4 |
| N-methylnicotinamide | 71.4 | -0.5 | 14.5 |
| Olmesartan | 60.4 | 37.0 | -7.9 |
| Ouabain | 50.5 | 20.5 | -24.1 |
| Piroxicam | 16.2 | 23.5 | 68.3 |
| Pravastatin | 52.2 | 5.4 | 36.8 |
| Progesterone | 63.9 | 33.4 | -362.6 |
| Quinine | 54.0 | -1.3 | -22.9 |
| Rosiglitazone | 79.1 | -32.1 | 10.0 |
| Saquinavir | 63.8 | 6.6 | 11.5 |
| Simvastatin | 73.1 | 29.3 | 47.4 |
| Spironolactone | 88.3 | -34.2 | -82.0 |
| Tetracycline | 22.1 | 29.5 | 51.1 |
| Valproic acid | 25.2 | 9.6 | 61.9 |
| Valsartan | 62.2 | 33.7 | 12.3 |
| Vincristine | 3.9 | 59.8 | 13.0 |
| 1-methyl-4-phenyl pyridinium | 14.2 | 10.8 | 2.4 |
| Acarbose | 14.9 | 17.7 | 0.1 |
| Aciclovir | 2.3 | -2.0 | -15.2 |
| Allopurinol | -35.6 | 10.8 | 8.4 |
| Amantadine | 26.4 | 10.0 | 17.0 |
| Amitriptyline | 10.5 | 22.0 | -8.9 |
| Amodiaquine | 24.2 | 28.2 | 43.4 |
| Atenolol | 20.6 | 7.1 | 6.8 |
| Atomoxetine | -7.5 | 31.6 | 16.3 |

| | | | |
|---|---|---|---|
| Berberine | 32.0 | 19.4 | 18.5 |
| Bestatin | 10.9 | 31.8 | 20.7 |
| Bufuralol | -13.9 | 23.8 | -7.0 |
| Bupropion | 8.5 | 7.2 | 17.0 |
| Buspirone | 33.3 | -5.6 | -13.9 |
| Caffeine | -1.8 | 22.1 | 3.2 |
| Captopril | 18.6 | 4.0 | 39.1 |
| Carbamazepine | 13.3 | 23.2 | 17.3 |
| Carnitine | 36.6 | -9.3 | 15.5 |
| Cefadroxil | 9.4 | 21.0 | 3.5 |
| Cefamandole | 30.2 | 16.5 | 2.7 |
| Celecoxib | -3.3 | 22.3 | -69.0 |
| Cetirizine | 35.7 | 40.3 | 29.9 |
| Chelerythrine | -0.7 | 24.9 | 13.1 |
| Chloroquine | -1.4 | -7.2 | 23.5 |
| Chlorpromazine | 27.1 | 22.8 | 4.3 |
| Chlorprothixene | 24.0 | -6.9 | 26.6 |
| Chlorzoxazone | 20.1 | -1.0 | 12.0 |
| Cholic acid | 41.6 | 40.8 | 20.2 |
| Cimetidine | 40.3 | 24.8 | -6.2 |
| Clotrimazole | 32.3 | -120.4 | -37.1 |
| Colchicine | 45.2 | -3.9 | 24.4 |
| Coumarin | 7.6 | 38.2 | 9.3 |
| Daidzein | 35.0 | 0.7 | 32.0 |
| Desipramine | 25.9 | 3.2 | -4.4 |
| Dexamethasone | 10.7 | 34.2 | -12.5 |
| Dextromethorphan | 36.3 | -21.9 | 15.9 |
| Digoxin | 36.0 | -4.7 | 32.1 |
| Diltiazem | 3.2 | 4.1 | -9.1 |
| Disopyramide | 47.5 | 8.6 | 18.5 |
| Disulfiram | 8.5 | -3.8 | 15.5 |
| Dofetilide | 1.0 | 19.9 | 18.1 |
| Doxazosin | 28.8 | -17.3 | 45.8 |
| Doxorubicin | -5.8 | 26.6 | 49.9 |
| Efavirenz | 40.9 | 18.2 | 43.2 |
| Eletriptan | 31.7 | 2.8 | 0.4 |
| Emtricitabine | 6.4 | 17.1 | 20.7 |
| Enalapril | -7.5 | 4.1 | 19.7 |
| Etoposide | 42.2 | 8.5 | 6.8 |
| Felodipine | 36.6 | 18.4 | 21.4 |
| Fendiline | 15.0 | 20.4 | -118.7 |
| Fenofibrate | 5.4 | -10.8 | 34.2 |
| Fentanyl | -53.3 | 22.6 | 11.7 |

| | | | |
|---|---|---|---|
| Fexofenadine | 28.9 | 6.0 | -5.8 |
| Fluconazole | 7.4 | 7.5 | 34.1 |
| Fluoxetine | 21.8 | 14.3 | 22.6 |
| Flupenthixol | 30.2 | 13.5 | 9.5 |
| Fluvoxamine | 43.8 | 29.2 | 39.0 |
| Furafylline | -14.7 | 0.7 | 18.1 |
| Furosemide | 23.4 | 22.3 | 34.5 |
| Glipizide | 1.1 | 25.2 | 22.0 |
| Glycyl proline | -8.4 | 9.6 | 24.1 |
| Hygromycin | 32.8 | -4.2 | 31.5 |
| Ibuprofen | 47.3 | -6.6 | 13.5 |
| Imipramine | 26.9 | -2.4 | 16.9 |
| Irinotecan | 40.5 | 26.9 | 16.3 |
| Isoniazid | 29.9 | 18.7 | 23.8 |
| Isradipine | 47.5 | 7.7 | 12.0 |
| Lamotrigine | 24.9 | 14.5 | 13.6 |
| Lansoprazole | 31.4 | 23.3 | -75.7 |
| Lisinopril | 4.6 | 9.0 | 19.2 |
| Loperamide | 32.5 | 32.4 | 19.6 |
| Loratadine | 43.2 | 45.0 | 28.1 |
| Mephenytoin | -5.7 | -0.6 | 32.9 |
| Metformin | -3.3 | 19.6 | -4.5 |
| Methotrexate | 27.4 | 17.2 | -13.4 |
| Methoxsalen | 38.1 | 8.9 | -1.6 |
| Metoprolol | 28.1 | 19.8 | 30.5 |
| Midazolam | 32.9 | 6.7 | 22.7 |
| Moclobemide | 1.7 | -3.3 | 27.5 |
| Naringenin | 34.8 | -57.2 | 32.4 |
| Naringin | 34.0 | 37.7 | 47.1 |
| Nicotine | 6.9 | 11.0 | -16.5 |
| Nitrofurantoin | 23.3 | -1.5 | 23.1 |
| N-methylpyridinium | 27.9 | -2.3 | 12.8 |
| N-methyl-quinidine | -6.1 | 21.0 | -10.3 |
| Nootkatone | 25.6 | -3.2 | -72.2 |
| Ofloxacin | -8.2 | 24.5 | 18.4 |
| Omeprazole | 15.9 | 29.5 | 11.3 |
| Ondansetron | 4.8 | -3.6 | -8.7 |
| P-aminohippuric acid | 30.1 | 23.7 | 27.4 |
| Pantoprazole | 15.2 | 23.4 | 17.3 |
| Paroxetine | 33.0 | 12.7 | 37.5 |
| Penicillin G | 8.8 | 40.9 | 7.6 |
| Phalloidin | 26.1 | 36.2 | 24.1 |
| Phenacetin | 34.2 | -0.9 | -0.1 |

| | | | |
|---|---|---|---|
| Phenformin | 34.5 | -2.3 | 16.7 |
| Phenobarbital | -3.9 | 7.3 | 11.0 |
| Phenylbutazone | 25.1 | 17.0 | 15.1 |
| Phenylethyl isothiocyanate | -16.4 | 12.3 | -32.0 |
| Phenytoin | 25.4 | 20.2 | 11.1 |
| Pilsicainide | 11.2 | -16.8 | 10.8 |
| Pindolol | -24.7 | 24.5 | -10.8 |
| Pioglitazone | 21.9 | 5.6 | -9.9 |
| Prazosin | 25.1 | 36.1 | 1.0 |
| Prednisolone | 2.3 | 9.6 | 27.6 |
| Probenecid | 35.2 | 22.0 | 26.4 |
| Procainamide | 21.6 | 3.4 | 17.8 |
| Propranolol | 8.9 | 12.8 | 10.1 |
| Quinidine | 43.3 | 37.2 | -10.0 |
| Ranolazine | 4.3 | -14.9 | 41.6 |
| Sanguinarine | -0.2 | 2.9 | 15.0 |
| Sildenafil | 22.2 | 30.8 | 45.9 |
| Sotalol | 46.4 | 6.6 | 19.0 |
| Sulfaphenazole | 16.9 | 0.7 | 17.1 |
| Tamoxifen | 25.6 | 28.5 | 11.0 |
| Tenofovir | 18.4 | 33.6 | 23.3 |
| Terfenadine | 42.7 | 33.3 | 3.7 |
| Testosterone | 33.2 | 37.4 | -175.1 |
| Tetraethylammonium | 25.9 | 4.9 | 6.7 |
| Theofylline | 0.8 | 2.5 | -25.5 |
| Thioridazine | 26.1 | 25.2 | 21.5 |
| Thiotepa | -40.3 | 0.6 | 16.2 |
| Ticlopidine | 19.4 | 0.2 | 23.0 |
| Tolbutamide | 9.1 | 5.8 | 15.5 |
| Topotecan | 36.7 | 17.3 | -39.2 |
| Tranylcypromine | -9.5 | -9.3 | 15.5 |
| Triazolam | 45.7 | 18.7 | -20.5 |
| Trimethoprim | 24.7 | 18.8 | 35.1 |
| Valaciclovir | 8.3 | 6.6 | -7.3 |
| Varenicline | 24.7 | 0.8 | 27.3 |
| Warfarin | 27.8 | -3.2 | 31.6 |
| Verapamil | 40.3 | -9.0 | -62.6 |
| Zidovudine | 9.5 | 7.1 | 4.7 |

**D.** Inhibition assay conditions from Karlgren et al. (2012).

| Transporter | Test Concentration | Probe Substrate | Probe Concentration |
|---|---|---|---|
| OATP1B1 | 20 µM | E17βG | 0.52 µM |
| OATP1B3 | 20 µM | E17βG | 1.04 µM |
| OATP2B1 | 20 µM | E3S | 1.02 µM |

**Supplemental Table S2: Structural-metric inventory by model**

**A.** AlphaFold3

| Metric Family | Quantity | Feature Names |
|---|---|---|
| Interface Confidence | 18 | struct_chain_iptm_0 |
| | | struct_chain_iptm_1 |
| | | struct_chain_iptm_max |
| | | struct_chain_iptm_mean |
| | | struct_chain_iptm_min |
| | | struct_chain_iptm_std |
| | | struct_chain_pair_iptm_0_0 |
| | | struct_chain_pair_iptm_0_1 |
| | | struct_chain_pair_iptm_1_0 |
| | | struct_chain_pair_iptm_1_1 |
| | | struct_chain_pair_iptm_max |
| | | struct_chain_pair_iptm_max_offdiag |
| | | struct_chain_pair_iptm_mean |
| | | struct_chain_pair_iptm_mean_offdiag |
| | | struct_chain_pair_iptm_min |
| | | struct_chain_pair_iptm_min_offdiag |
| | | struct_chain_pair_iptm_std |
| | | struct_iptm |
| Fold / Local Confidence | 7 | struct_chain_ptm_0 |
| | | struct_chain_ptm_1 |
| | | struct_chain_ptm_max |
| | | struct_chain_ptm_mean |
| | | struct_chain_ptm_min |
| | | struct_chain_ptm_std |
| | | struct_ptm |
| Predicted Error | 11 | struct_chain_pair_pae_min_0_0 |
| | | struct_chain_pair_pae_min_0_1 |
| | | struct_chain_pair_pae_min_1_0 |
| | | struct_chain_pair_pae_min_1_1 |
| | | struct_chain_pair_pae_min_max |
| | | struct_chain_pair_pae_min_max_offdiag |
| | | struct_chain_pair_pae_min_mean |
| | | struct_chain_pair_pae_min_mean_offdiag |
| | | struct_chain_pair_pae_min_min |
| | | struct_chain_pair_pae_min_min_offdiag |
| | | struct_chain_pair_pae_min_std |
| Ranking / Aggregate | 1 | struct_ranking_score |
| Clashes / Disorder | 2 | struct_fraction_disordered |

| | | |
|---|---|---|
| | | struct_has_clash |

**B.** Boltz-2

| Metric Family | Quantity | Feature Names |
|---|---|---|
| Interface Confidence | 8 | struct_complex_iplddt |
| | | struct_iptm |
| | | struct_ligand_iptm |
| | | struct_pair_chains_iptm_0_0 |
| | | struct_pair_chains_iptm_0_1 |
| | | struct_pair_chains_iptm_1_0 |
| | | struct_pair_chains_iptm_1_1 |
| | | struct_protein_iptm |
| Fold / Local Confidence | 4 | struct_chains_ptm_0 |
| | | struct_chains_ptm_1 |
| | | struct_complex_plddt |
| | | struct_ptm |
| Predicted Error | 2 | struct_complex_ipde |
| | | struct_complex_pde |
| Ranking / Aggregate | 1 | struct_confidence_score |
| Clashes / Disorder | 0 | - |

**C.** Chai-1

| Metric Family | Quantity | Feature Names |
|---|---|---|
| Interface Confidence | 12 | struct_iptm |
| | | struct_per_chain_pair_iptm_0_0_0 |
| | | struct_per_chain_pair_iptm_0_0_1 |
| | | struct_per_chain_pair_iptm_0_1_0 |
| | | struct_per_chain_pair_iptm_0_1_1 |
| | | struct_per_chain_pair_iptm_max |
| | | struct_per_chain_pair_iptm_max_offdiag |
| | | struct_per_chain_pair_iptm_mean |
| | | struct_per_chain_pair_iptm_mean_offdiag |
| | | struct_per_chain_pair_iptm_min |
| | | struct_per_chain_pair_iptm_min_offdiag |
| | | struct_per_chain_pair_iptm_std |
| Fold / Local Confidence | 7 | struct_per_chain_ptm_0_0 |
| | | struct_per_chain_ptm_0_1 |
| | | struct_per_chain_ptm_max |
| | | struct_per_chain_ptm_mean |
| | | struct_per_chain_ptm_min |

| | | |
|---|---|---|
| | | struct_per_chain_ptm_std |
| | | struct_ptm |
| Predicted Error | 0 | - |
| Ranking / Aggregate | 1 | struct_aggregate_score |
| Clashes / Disorder | 11 | struct_chain_chain_clashes_0_0_0 |
| | | struct_chain_chain_clashes_0_0_1 |
| | | struct_chain_chain_clashes_0_1_0 |
| | | struct_chain_chain_clashes_0_1_1 |
| | | struct_chain_chain_clashes_max |
| | | struct_chain_chain_clashes_max_offdiag |
| | | struct_chain_chain_clashes_mean |
| | | struct_chain_chain_clashes_mean_offdiag |
| | | struct_chain_chain_clashes_min |
| | | struct_chain_chain_clashes_min_offdiag |
| | | struct_chain_chain_clashes_std |

**D.** Protenix

| Metric Family | Quantity | Feature Names |
|---|---|---|
| Interface Confidence | 29 | struct_chain_iptm_0 |
| | | struct_chain_iptm_1 |
| | | struct_chain_iptm_max |
| | | struct_chain_iptm_mean |
| | | struct_chain_iptm_min |
| | | struct_chain_iptm_std |
| | | struct_chain_pair_iptm_0_0 |
| | | struct_chain_pair_iptm_0_1 |
| | | struct_chain_pair_iptm_1_0 |
| | | struct_chain_pair_iptm_1_1 |
| | | struct_chain_pair_iptm_global_0_0 |
| | | struct_chain_pair_iptm_global_0_1 |
| | | struct_chain_pair_iptm_global_1_0 |
| | | struct_chain_pair_iptm_global_1_1 |
| | | struct_chain_pair_iptm_global_max |
| | | struct_chain_pair_iptm_global_max_offdiag |
| | | struct_chain_pair_iptm_global_mean |
| | | struct_chain_pair_iptm_global_mean_offdiag |
| | | struct_chain_pair_iptm_global_min |
| | | struct_chain_pair_iptm_global_min_offdiag |
| | | struct_chain_pair_iptm_global_std |
| | | struct_chain_pair_iptm_max |
| | | struct_chain_pair_iptm_max_offdiag |

| | | |
|---|---|---|
| | | struct_chain_pair_iptm_mean |
| | | struct_chain_pair_iptm_mean_offdiag |
| | | struct_chain_pair_iptm_min |
| | | struct_chain_pair_iptm_min_offdiag |
| | | struct_chain_pair_iptm_std |
| | | struct_iptm |
| Fold / Local Confidence | 25 | struct_chain_pair_plddt_0_0 |
| | | struct_chain_pair_plddt_0_1 |
| | | struct_chain_pair_plddt_1_0 |
| | | struct_chain_pair_plddt_1_1 |
| | | struct_chain_pair_plddt_max |
| | | struct_chain_pair_plddt_max_offdiag |
| | | struct_chain_pair_plddt_mean |
| | | struct_chain_pair_plddt_mean_offdiag |
| | | struct_chain_pair_plddt_min |
| | | struct_chain_pair_plddt_min_offdiag |
| | | struct_chain_pair_plddt_std |
| | | struct_chain_plddt_0 |
| | | struct_chain_plddt_1 |
| | | struct_chain_plddt_max |
| | | struct_chain_plddt_mean |
| | | struct_chain_plddt_min |
| | | struct_chain_plddt_std |
| | | struct_chain_ptm_0 |
| | | struct_chain_ptm_1 |
| | | struct_chain_ptm_max |
| | | struct_chain_ptm_mean |
| | | struct_chain_ptm_min |
| | | struct_chain_ptm_std |
| | | struct_plddt |
| | | struct_ptm |
| Predicted Error | 18 | struct_chain_gdpe_0 |
| | | struct_chain_gdpe_1 |
| | | struct_chain_gdpe_max |
| | | struct_chain_gdpe_mean |
| | | struct_chain_gdpe_min |
| | | struct_chain_gdpe_std |
| | | struct_chain_pair_gdpe_0_0 |
| | | struct_chain_pair_gdpe_0_1 |
| | | struct_chain_pair_gdpe_1_0 |
| | | struct_chain_pair_gdpe_1_1 |
| | | struct_chain_pair_gdpe_max |
| | | struct_chain_pair_gdpe_max_offdiag |

| | | |
|---|---|---|
| | | struct_chain_pair_gdpe_mean |
| | | struct_chain_pair_gdpe_mean_offdiag |
| | | struct_chain_pair_gdpe_min |
| | | struct_chain_pair_gdpe_min_offdiag |
| | | struct_chain_pair_gdpe_std |
| | | struct_gdpe |
| Ranking Aggregate | 1 | struct_ranking score |
| Clashes / Disorder | 1 | struct_disorder |

**Supplemental Table S3: Structural descriptors selected by optimized single-metric classifiers.** For each protein–ligand structure-prediction pipeline and OATP isoform, the individual structural descriptor and L2 regularization strength were selected independently within each outer training partition using inner cross-validation optimized for average precision (AP). The same held-out predictions from these optimized single-descriptor classifiers were used to calculate both AP and ROC-AUC in Figure 2A–B. Descriptor names correspond to the original model-derived feature names.

| Structural Model | OATP Isoform | Selected Metric | Interpretation |
|---|---|---|---|
| AlphaFold 3 | OATP1B1 | chain_pTM | Chain-level TM confidence |
| AlphaFold 3 | OATP1B3 | chain_pair_ipTM | Chain-pair interface confidence |
| AlphaFold 3 | OATP2B1 | chain_pTM | Chain-level TM confidence |
| Boltz-2 | OATP1B1 | complex_pLDDT | Local complex confidence |
| Boltz-2 | OATP1B3 | chains_pTM_1 | TM score for indexed chain 1 |
| Boltz-2 | OATP2B1 | chains_pTM_1 | TM score for indexed chain 1 |
| Chai-1 | OATP1B1 | pair_ipTM_0_0_1 | Chain pair interface confidence |
| Chai-1 | OATP1B3 | pair_ipTM_0_0_1 | Chain pair interface confidence |
| Chai-1 | OATP2B1 | pair_ipTM_min | Min. chain interface confidence |
| Protenix | OATP1B1 | chain_pTM | Chain-level TM confidence |
| Protenix | OATP1B3 | chain_pTM | Chain-level TM confidence |
| Protenix | OATP2B1 | GDPE | Global predicted distance error |
